# Reduced physiological reserve and a blunted response to acute exercise in adults with cerebral palsy

**DOI:** 10.64898/2026.10.11.758338

**Authors:** Linnéa Corell, Oscar Horwath, Sebastian Edman, Stefan M Reitzner, William Hangasjärvi, Guadalupe Meza, Jenny Castaings, Annika Kruse, Julia Starck, Niklas Valtonen, Emma Hjalmarsson, Ola Kvist, Daniele A. Cardinale, Eva Pontén, Peder Sörensson, Rodrigo Fernandez-Gonzalo, Sudarshan Dayanidhi, Mark D Peterson, Jessica Norrbom, Ferdinand von Walden

## Abstract

Cerebral palsy (CP) is an early-onset neurodevelopmental condition associated with metabolic impairment and an increased risk of non-communicable diseases across the lifespan. However, the basis of these changes remains largely unknown. This study provides a comprehensive cardiometabolic and physical function profile of 23 adults with CP relative to 33 typically developing (TD) adolescents and 61 TD adults. Compared to TD individuals, adults with CP exhibited reduced cardiorespiratory fitness, smaller cardiac dimensions, and lower muscle mass. Circulating metabolomic and proteomic profiling identified a distinct molecular signature in CP, including altered tricarboxylic acid (TCA) cycle metabolites, and elevated myostatin, matrix-related proteins and pro-inflammatory cytokines. We further tested the capacity for physiological and molecular adaptation using an acute bout of endurance exercise. In response to exercise, individuals with CP exhibited an attenuated response compared to TD groups, characterized by a blunted lactate response and lower venous oxygen levels. TD participants also increased canonical exercise-responsive factors, including NADK, IL-8, ANGPTL1 and GDF15, whereas these responses were attenuated in CP. These novel findings define a distinct cardiometabolic profile in adults with CP at rest and reveal an attenuated molecular response to exercise. These data carry implications for targeted interventions and prevention strategies across the lifespan.

## INTRODUCTION

Cerebral palsy (CP) is the most common cause of early-onset neurodevelopmental physical disability, affecting ∼1.6-3.4 per 1000 live births and an estimated 17 million people worldwide (1–3). It results from injury or malformation of the developing brain and is characterized by permanent motor impairments of varying type and severity, with gross motor function classified according to the Gross Motor Function Classification System (GMFCS) (4, 5). These impairments are typically accompanied by pain, fatigue, and reduced mobility, contributing to limitations in the amount of daily physical activity, which become more pronounced at higher GMFCS levels (6–8). While the initial brain injury is static, CP is associated with progressive secondary health complications that accumulate across the lifespan (9, 10), including a premature decline in motor function (11). Adults with CP also face a markedly increased risk of several non-communicable diseases, including cardiometabolic comorbidities (6, 12–15). Despite this growing recognition of adverse health outcomes, the physiological mechanisms underlying premature functional decline and elevated disease risk in adulthood remain poorly understood.

Cardiometabolic disease risk is elevated among adults with CP, who exhibit a high prevalence of established risk factors, including hypertension, dyslipidemia, and excess adiposity (12, 16–18). This adverse cardiometabolic profile is accompanied by consistently lower cardiorespiratory fitness (CRF), with peak oxygen uptake (VO_2_ peak) typically 23-40% lower than that of age-matched typically developing (TD) individuals (19–21). Given that CRF is a robust indicator of cardiovascular function and an independent predictor of mortality (22), this reduction carries important clinical implications. The reduced VO_2_ peak observed in adults with CP likely reflects the combined influence of cardiorespiratory, muscular, and systemic limitations (23, 24). Moreover, the well-documented low levels of physical activity and/or excessive sedentariness observed in this population may further contribute to reduced CRF, compounding their already elevated cardiometabolic risk (7).

Regular physical activity and exercise are fundamental to health and cardiovascular risk reduction (25). These benefits arise through repeated bouts of exercise that transiently induce coordinated physiological and molecular responses across multiple organ systems, ultimately driving long-term adaptation (26–28). However, functional limitations associated with CP may constrain exercise frequency, volume and intensity, potentially attenuating exercise-induced adaptations (29, 30). In addition to reduced exercise stimulus, intrinsic factors may further limit the adaptive response. For instance, skeletal muscle in individuals with CP exhibits features that may affect its capacity to adapt to increased physiological demands, including shifts in muscle fiber type (31), reduced satellite cell content, and changes in pathways involved in muscle mass regulation (32). Alterations in circulating mediators of exercise adaptation, including immune cell populations (33), extracellular vesicles (34), and metabolites (35), further suggest that the systemic signals necessary for exercise adaptations are altered in this population. These observations raise the possibility that the capacity to mount a robust and coordinated response to exercise differs in CP. While we have previously provided insight into selected aspects of this response, including direct comparisons with individuals without CP (19, 33), a comprehensive characterization of the acute systemic response remains lacking.

Despite recent advances, cardiometabolic physiology in adults with CP remains poorly understood. To address this, we performed physiological profiling across several domains in adults with CP, encompassing cardiac characteristics and function, skeletal muscle size, and circulating molecular signatures. These data were compared to a large reference cohort of TD adults as well as TD adolescents, to enable comparisons across age and body size. Compared with TD adults, adults with CP exhibited reduced aerobic capacity, structurally smaller hearts, and a systemic milieu enriched in pro-inflammatory cytokines, matrix-related proteins and negative regulators of muscle mass. These features indicate a basal physiological state that may constrain cardiovascular and skeletal muscle function. Furthermore, in response to an acute bout of endurance exercise, adults with CP demonstrated a limited response to physiological stress, with blunted metabolic and blood gas responses and attenuated changes in the plasma proteome. Our findings suggest that CP might be associated with a constrained physiological reserve that may contribute to elevated cardiometabolic vulnerability. These findings also highlight the importance of understanding how exercise capacity and adaptive responses may be optimized in individuals with CP and support the need to develop targeted intervention strategies in this population.

## RESULTS

### Participant characteristics and maximal cardiorespiratory capacity

We studied 23 adults with CP (18-44 yrs) alongside two control groups: 33 TD adolescents (13-17 yrs) and 61 TD adults (18-50 yrs). The inclusion of both control groups was intended to provide complementary reference populations, with TD adults serving as an adult comparison group and TD adolescents providing a comparison for body size, as adults with CP often exhibit reduced stature and altered growth trajectories (36). In the resulting sample, the age of adults with CP fell between that of the two TD groups (median age 23 vs. 15 and 39 yrs, respectively), but more closely resembled TD adolescents in terms of body mass, height, and BMI (Table 1). The CP cohort was clinically heterogeneous, spanning GMFCS levels II-IV. The cohort consisted predominantly of spastic (52%) and dyskinetic (35%) subtypes, with the remaining participants classified as ataxic (9%). Individuals with CP demonstrated markedly reduced CRF. During maximal treadmill exercise to task failure (Table 1), their absolute VO_2_ peak was 36-40% lower than in both TD groups and remained 20-30% lower when expressed relative to body mass. Anthropometric data, 6-Minute Frame Running Test (6-MFRT) performance and clinical characteristics are also provided in Table 1. A statistical summary of all group comparisons is provided in Table S1.

**Table 1.**
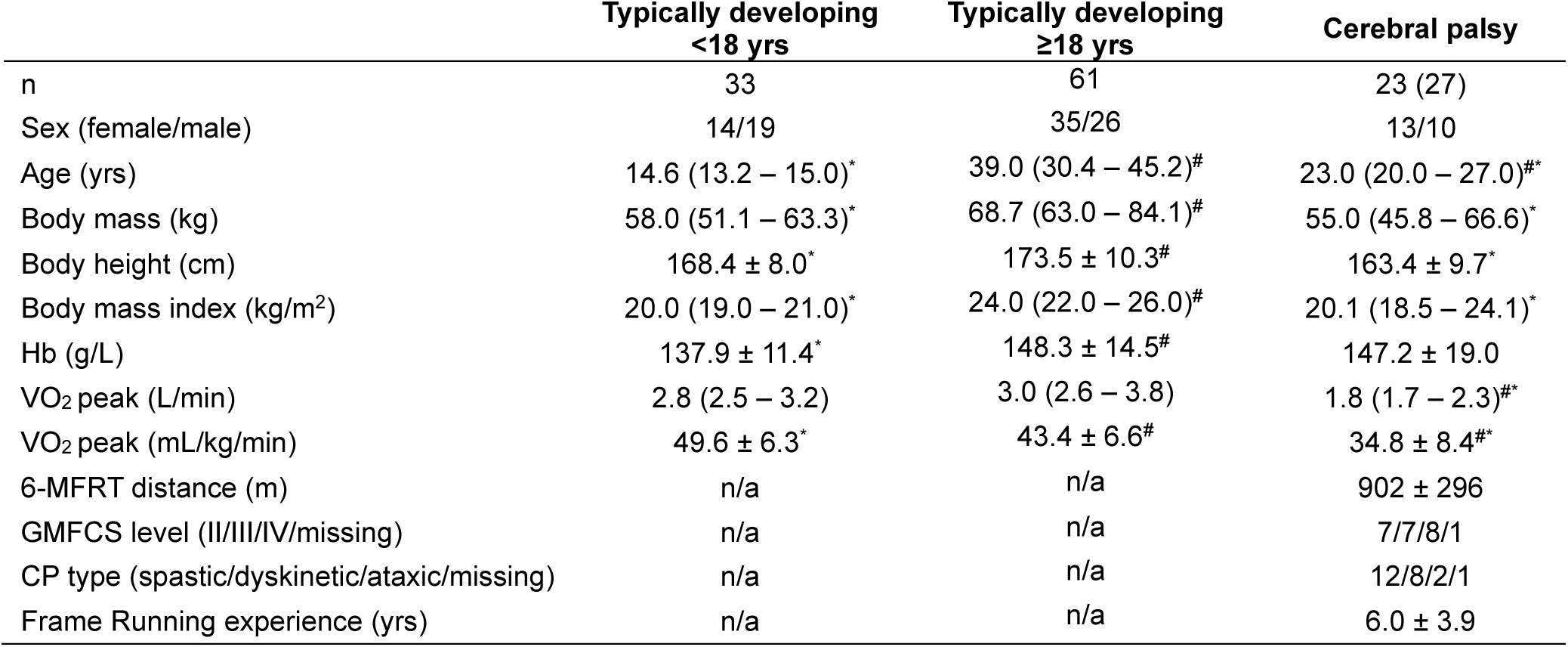
Demographic, anthropometric, and clinical characteristics of CP and TD individuals. Data are presented as mean ± SD for normally distributed variables or median (IQR) for non-normally distributed variables. For the CP group, n is given as unique individuals (observations). Twenty-three adults with CP contributed 27 observations, as two individuals were assessed twice and one individual three times (see Methods). Sex, GMFCS level, and CP type are reported for unique individuals (n=23), whereas all other characteristics are reported per observation (see Table S1, and Table S3). A valid maximal exercise test was obtained in 33 TD adolescents, 61 TD adults, and 10 of the 12 adults with CP who attempted the test. The remaining two did not fulfill the criteria for a valid physiological maximal exercise response (see Methods). A valid 6-MFRT was obtained in 22 of the 27 CP observations. In one case the criteria for a valid maximal physiological exercise response were not met, and in four the test was not performed or data were missing. Between-group differences were assessed using one-way ANOVA followed by Tukey’s multiple comparisons test for normally distributed variables, or Kruskal-Wallis test followed by Dunn’s multiple comparisons test for non-normally distributed variables. # indicates a significant difference from TD adolescents, * indicates a significant difference from TD adults. Statistical significance was defined as adjusted p < 0.05. Abbreviations: CP, cerebral palsy; GMFCS, Gross Motor Function Classification System; Hb, hemoglobin; IQR, interquartile range; n/a, not applicable; SD, standard deviation; VO_2_ peak, peak oxygen uptake.

### Structurally smaller hearts, but no overt signs of cardiac dysfunction in adults with CP

Resting cardiac structure and function were assessed by electrocardiogram (ECG) and echocardiography (Figure 1A). Resting blood pressure was within the normotensive range across all groups and ECG conduction and repolarization parameters did not differ between CP and TD participants (Table S2). However, resting heart rate was higher in CP than in TD adults, with a similar trend when compared to TD adolescents (median differences of 14 and 10 beats per minute, respectively, *p=*0.008 vs TD adults*, p*=0.052 vs TD adolescents; Table S2).

**Figure 1.**
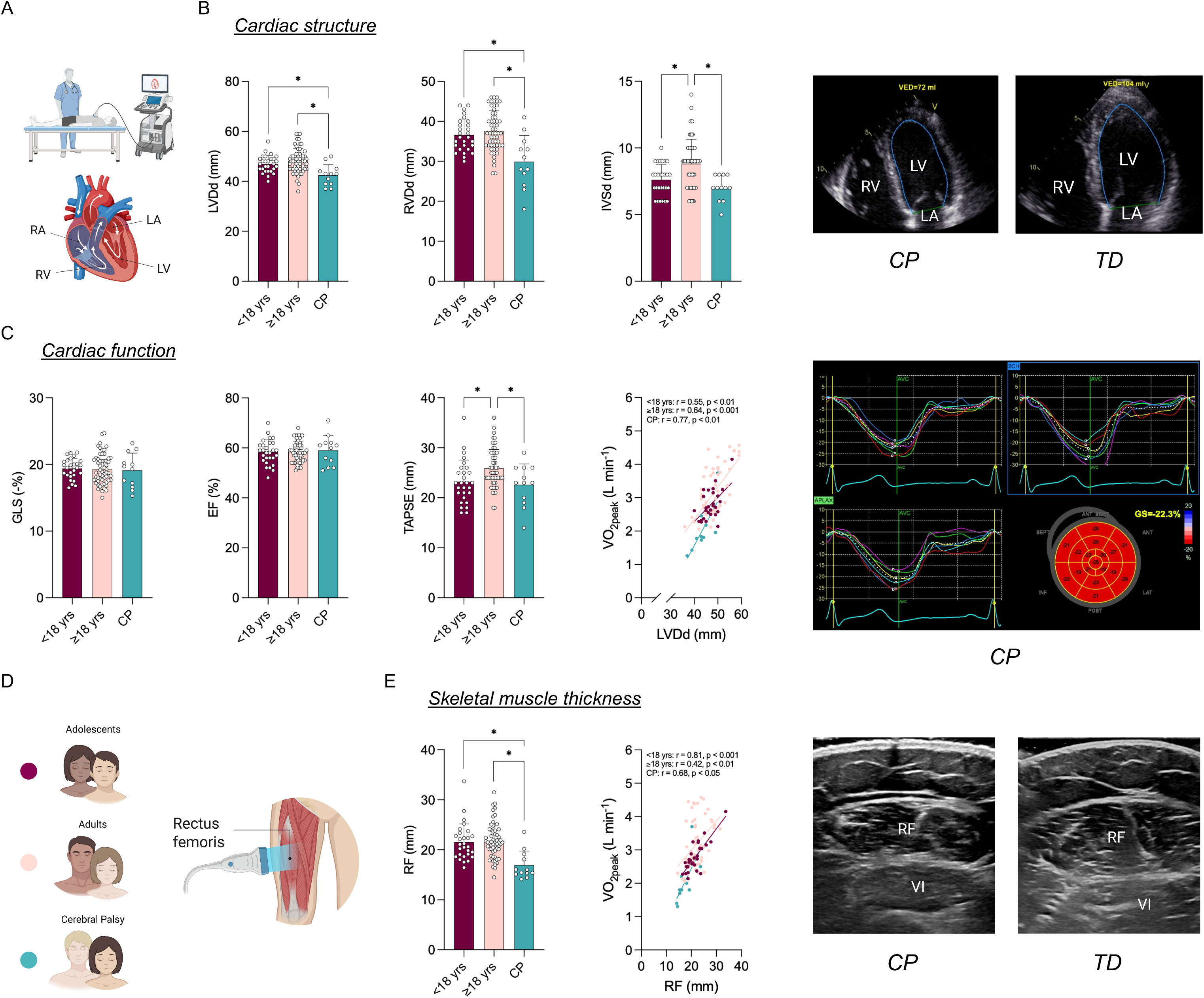
Cardiac structure and function, and skeletal muscle thickness in CP and TD individuals. (A) Schematic overview of the transthoracic echocardiographic assessment and cardiac anatomy. (B) Left ventricular diameter at end-diastole (LVDd), right ventricular diameter at end-diastole (RVDd), and interventricular septal thickness at end-diastole (IVSd) in TD adolescents, TD adults, and individuals with CP. Representative apical four-chamber echocardiographic images from a participant with CP and a TD participant. (C) Global longitudinal strain (GLS), ejection fraction (EF), and tricuspid annular plane systolic excursion (TAPSE) in the three groups. Correlation between peak oxygen uptake (VO_2_ peak) and LVDd within each group, with Pearson correlation coefficients (r) and p values indicated. Representative plot of global strain from a participant with CP. (D) Schematic overview of the study groups (TD adolescents, TD adults, individuals with CP) and B-mode ultrasonographic assessment of m. rectus femoris (RF) muscle thickness. (E) RF thickness in the three groups. Correlation between peak oxygen uptake (VO_2_ peak) and RF thickness within each group, with Pearson correlation coefficients (r) and p values indicated. Representative ultrasound images from a participant with CP and a TD participant. Sample size: CP *n* = 12, TD adolescents *n* = 28, and TD adults *n* = 58. Significant differences between groups were assessed using one-way ANOVA with Tukey’s post hoc test, or Kruskal-Wallis with Dunn’s test for non-normally distributed data. * indicates adjusted p < 0.05. Data are presented as mean ± SD for each group.

Furthermore, echocardiographic analyses revealed smaller chamber dimensions in adults with CP. Both left and right ventricular end-diastolic diameters (LVDd and RVDd) were smaller in CP, with LVDd reduced by ∼10–12% and RVDd by ∼18–20%, compared with TD adolescents and adults (Figure 1B). When indexed to body surface area (LVDi and RVDi, Figure S1), these differences were attenuated. LVDi differed between the two TD groups, whereas adults with CP did not differ from either group. RVDi followed a similar pattern, differing between the two TD groups, with an additional difference between CP and TD adolescents but not between CP and TD adults. Interventricular septal thickness (IVSd) differed primarily between TD adults and the other groups, while CP and TD adolescents showed comparable values (Figure 1B). When indexed (IVSi, Figure S1), no differences were observed between the three groups.

Despite these structural differences, global longitudinal strain (GLS) and left ventricular ejection fraction (EF) were within the normal range and did not differ between groups, indicating preserved left ventricular systolic function in individuals with CP (Figure 1C). Tricuspid annular plane systolic excursion (TAPSE) was higher in TD adults than in both TD adolescents and CP, which did not differ from each other (Figure 1C). Other echocardiographic indices showed no group differences (S′ velocity, left atrial volume index), while diastolic parameters followed expected age-related patterns without a consistent deviation related to CP (Figure S1). The functional relevance of cardiac size was further explored by relating LVDd to aerobic capacity. LVDd correlated positively with VO_2_ peak across all three groups, with the CP cohort clustering at the lower end of both distributions, consistent with their lower aerobic capacity (Figure 1C).

Given the structural differences observed in the heart and its potential relevance to aerobic capacity, muscle thickness of *m. rectus femoris* was assessed as a proxy for skeletal muscle mass (Figure 1D). Muscle thickness was markedly lower in adults with CP compared to both TD adolescents and TD adults, by approximately 25% and 27%, respectively (Figure 1E). Similar to LVDd, rectus femoris thickness correlated positively with VO_2_ peak across all three groups, with the CP cohort again clustering toward the lower end of the respective distributions (Figure 1E), suggesting that reduced peripheral muscle mass may be a contributor to lower aerobic capacity in CP. To determine whether this peripheral deficit was related to the cardiac phenotype described above, we tested the relationship between rectus femoris thickness and LVDd across groups (Figure S1). A positive association was present in TD adolescents, but not in TD adults or in CP. Additionally, within the CP group, cardiac dimensions and function did not differ between participants classified as GMFCS II-III (primarily ambulatory) and those at GMFCS level IV (primarily using a wheelchair for mobility). Similarly, rectus femoris thickness did not differ between GMFCS levels (data not shown).

### CP is associated with a distinct basal plasma molecular signature, with subtle variations according to motor function severity

The systemic molecular profile was characterized using targeted liquid chromatography-mass spectrometry-based metabolomics on basal plasma samples, yielding quantification of 261 hydrophilic metabolites. Principal component analysis (PCA) showed substantial overlap between CP and TD adolescents and adults, with no clear separation and variance distributed across PC1/PC2 (Figure 2A). Subsequent differential analysis identified 13 metabolites (5.0% of the 261 quantified) that were differentially abundant in CP (increased or decreased). The tricarboxylic acid (TCA) cycle-associated metabolites 2-hydroxyglutarate (2-HG) and aconitic acid were reduced in CP compared with both TD groups (Figure 2B). In addition, alterations in tryptophan metabolism were observed in CP, with lower levels of 3-hydroxykynurenine and elevated levels of serotonin. Metabolites involved in one-carbon and methylation metabolism were also altered, including decreased S-adenosylmethionine and 5-methylthioadenosine (Figure S2).

**Figure 2.**
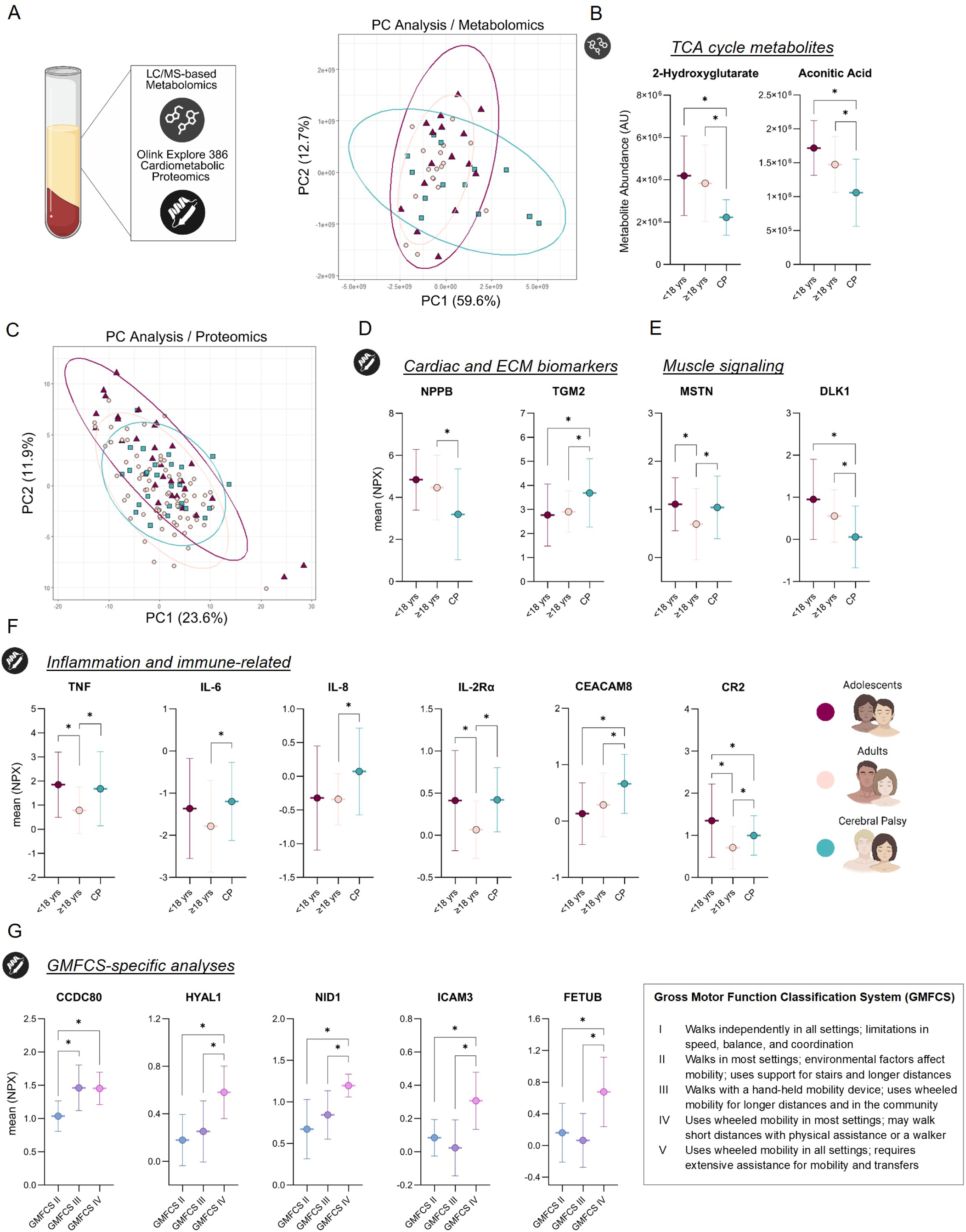
Basal circulating metabolite and protein profile in CP and TD individuals and GMFCS-specific protein analysis in CP. (A) Schematic overview of analytical panels and a principal component analysis (PCA) of basal plasma metabolites. Each symbol represents a plasma sample, colored by group (purple, TD adolescents; pink, TD adults; green, CP adults). (B) Metabolites associated with the TCA cycle. (C) PCA of basal plasma proteins. Each symbol represents a plasma sample, colored by group (purple, TD adolescents; pink, TD adults; green, CP adults). (D) Cardiac- and ECM-related proteins. (E) Skeletal muscle-related proteins. (F) Inflammation- and immune system-related proteins. (G) GMFCS-specific differences in circulating proteins among adults with CP, with definitions of GMFCS levels I-V provided in the accompanying information box. PCA was used to visualize the overall variation in basal plasma metabolite and protein profiles, with samples plotted according to their scores on PC1 and PC2. Sample size for metabolites/proteins: CP *n* = 13/27, TD adolescents *n* = 13/30, and TD adults *n* = 17/59. Significant differences in metabolite and protein levels between groups, as well as differences in protein levels across GMFCS levels, were assessed using one-way ANOVA. P values were adjusted for multiple comparisons using Tukey’s post-hoc test. * indicates adjusted p < 0.05. Data are presented as mean ± SD for each group.

Structural alterations in cardiac and skeletal muscle size, alongside the metabolomic profile in CP, prompted further investigation of circulating proteins associated with cardiometabolic and musculoskeletal regulation. Using a proximity extension assay (PEA)-based proteomics approach (Olink Explore cardiometabolic panel), we quantified 366 plasma proteins. Expected age-related changes were observed for proteins known to change with growth and maturation (37–39), such as collagen type I alpha 1 chain (COL1A1), aggrecan (ACAN), and neuronal pentraxin receptor (NPTXR), reflecting robust biological signals in the dataset (Figure S3).

At the global level, PCA showed substantial overlap between all three groups, with no clear separation and variance distributed across PC1/PC2 (Figure 2C). Despite this similarity, further analysis revealed a subset of 39 proteins (11% of the 366 quantified) that were altered in CP, indicating selective perturbations in circulating proteins. Within proteins linked to cardiovascular biology, natriuretic peptide B (NPPB), a cardiac stress-related peptide released from cardiomyocytes, was reduced in CP (Figure 2D). Further, proteins associated with extracellular matrix (ECM) organization were altered, most notably an elevation of transglutaminase 2 (TGM2), a key mediator of ECM remodeling and stabilization (40) (Figure 2D). A similar pattern of increased abundance was observed for other ECM-associated proteins, including COL1A1, pleiotrophin (PTN), and serpin family B member 5 (SERPINB5) (Figure S3), which extends previous findings in pediatric CP cohorts where this profile is a typical feature (41).

Among skeletal muscle-related proteins, adults with CP had elevated levels of myostatin (MSTN), a negative regulator of muscle mass, and reduced levels of delta-like protein 1 (DLK1), a key modulator of myostatin signaling (42). Both changes are consistent with reduced muscle mass (Figure 2E). These changes were accompanied by differences in inflammatory and immune-related proteins, with increased levels of tumor necrosis factor (TNF) and interleukin-6 (IL-6) in adults with CP compared with TD adults (Figure 2F). Several additional markers showed higher levels in adults with CP than in TD adults. However, these differences were less consistent when adults with CP were compared with TD adolescents, consistent with normal variation in inflammatory markers across developmental stages (43, 44). In addition, adults with CP exhibited lower circulating levels of leptin, fatty acid-binding protein 4 (FABP4), and serpin family A member 12 (SERPINA12), together with elevated levels of hexokinase 2 (HK2), indicating a distinct metabolic protein profile (Figure S3).

To further assess whether the plasma protein signature in adults with CP could be related to deconditioning associated with limited mobility, we examined associations between 6-MFRT performance and a selection of these proteins, including TGM2, IL-6, IL-8, GDF15, TNF and MSTN. However, this analysis suggested that functional capacity was unlikely to be a major determinant of the observed protein differences as no significant correlations were observed (Figure S4). We followed this up by identifying common and distinct plasma proteomic signatures associated with physical activity in published datasets. Across four studies, circulating plasma proteins were examined in relation to habitual physical activity in large population-based cohorts, using questionnaire-based and/or device-based measures of physical activity (45–48). From each study, we extracted proteins strongly associated with physical activity from the reported lists and figures. Where available, we prioritized proteins with replicated associations in independent cohorts. Proteins were classified according to the reported direction of association as positive or negative. From our CP cohort, we selected the 15 most differentially abundant proteins relative to TD adults and compared these with those associated with reduced physical activity in the published datasets (Figure S4). Of the 10 increased proteins, only MSTN and IL-6 overlapped with proteins associated with lower physical activity, while none of the five decreased proteins overlapped with proteins associated with higher physical activity, indicating limited concordance with physical activity-associated plasma protein signatures.

We next examined whether the degree of motor impairment was associated with differences in the basal plasma metabolite and protein profiles in CP. For plasma metabolites, the sample size precluded analysis of individual GMFCS levels, and we therefore compared GMFCS II+III (primarily ambulatory) versus IV (primarily using a wheelchair for mobility). Only four metabolites differed significantly between groups, with GMFCS IV having higher 3-phosphoserine and cystathionine and lower guanidinoacetate and histamine (Figure S2). For plasma proteins, the larger sample size allowed us to stratify the CP cohort into three categories based on GMFCS levels (II, III, and IV). This approach yielded five proteins displaying a consistent pattern across GMFCS categories, with higher abundance in those with greater motor impairment. ECM-associated proteins, including coiled-coil domain containing 80 (CCDC80), hyaluronidase 1 (HYAL1), and nidogen-1 (NID1), were increased in the higher GMFCS groups, potentially reflecting alterations in ECM organization associated with greater cardiorespiratory and/or muscle impairment. In addition, intercellular adhesion molecule 3 (ICAM3), a transmembrane glycoprotein primarily expressed on leukocytes, and fetuin-B (FETUB), a liver-derived factor associated with impaired glucose metabolism (49), showed increased abundance with higher motor impairment (Figure 2G).

### Reduced endurance capacity and cardiovascular response to exercise in adults with CP despite similar perceived exertion

Participants completed an acute bout of endurance exercise consisting of Frame Running for adults with CP or conventional running for TD participants, during which they were instructed to cover as much distance as possible over 45 minutes on an indoor running track (Figure 3A). Despite reporting similar perceived exertion throughout the exercise session, adults with CP covered 43% less distance than TD adults and 41% less than TD adolescents (Figure 3B). Heart rate intensity (%) was also lower in individuals with CP and adolescents compared with adults (median intensity 82.9%, 90.0% and 91.8%, respectively) (Figure 3B).

**Figure 3.**
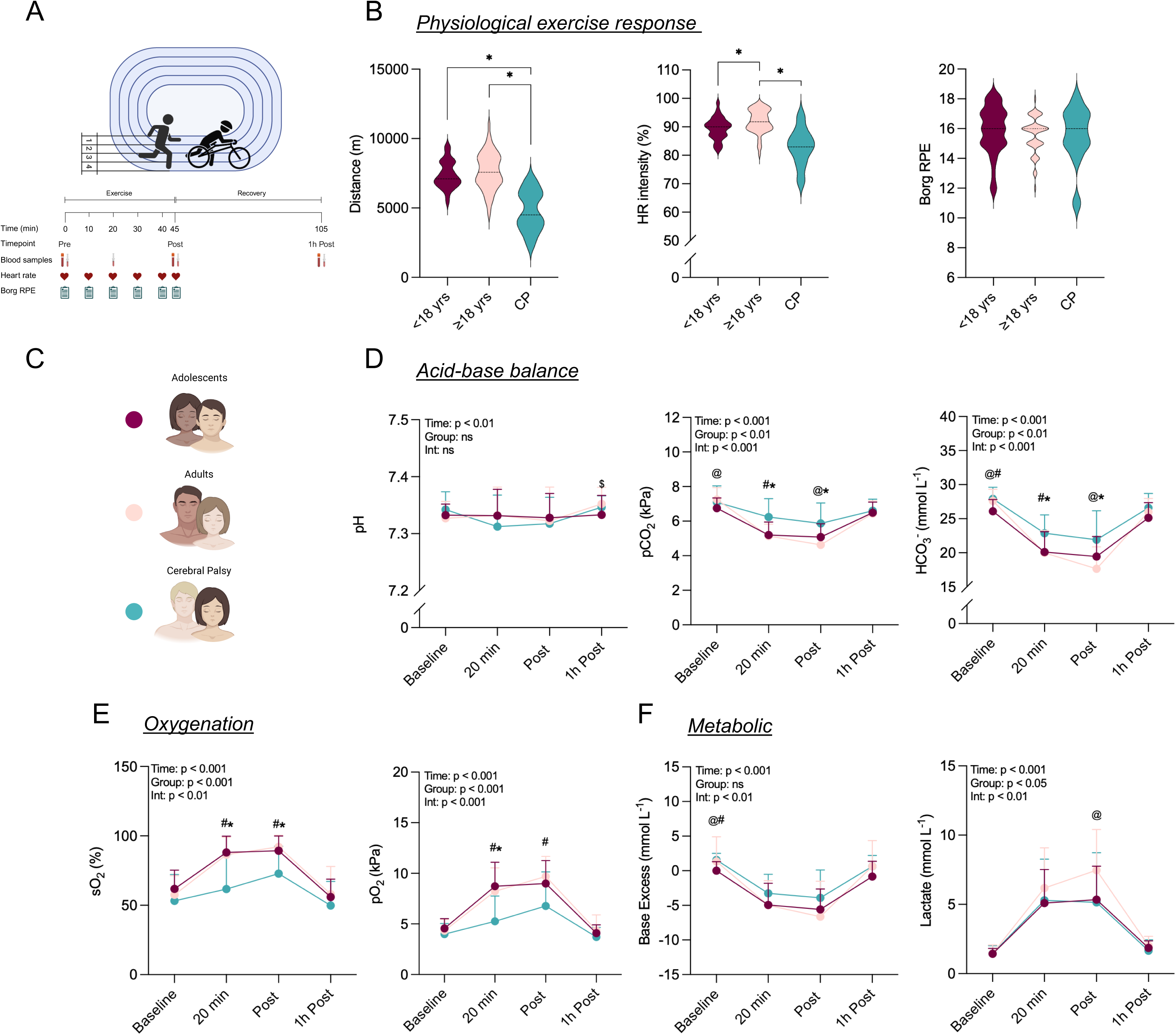
Physiological and acid-base response in CP and TD individuals to an acute bout of endurance exercise. (A) Schematic overview of the 45-minute exercise session protocol, including the exercise and recovery phases and the timing of venous blood sampling, heart rate monitoring, and Borg Rating of Perceived Exertion (RPE) assessments. (B) Distance covered, mean heart rate intensity (% of maximal heart rate), and weighted mean Borg RPE during the session in TD adolescents, TD adults, and individuals with CP. Violin plots display the distribution of individual values, with the median indicated by a dashed line. (C) Schematic overview of the study groups (TD adolescents, TD adults, individuals with CP). (D) pH, partial pressure of carbon dioxide (pCO_2_), and bicarbonate (HCO_3_^-^) at baseline, at 20 minutes, immediately post-exercise, and one-hour post-exercise. (E) Oxygen saturation (sO_2_), partial pressure of oxygen (pO_2_), (F) base excess and lactate at the same time points. Sample size: CP *n* = 12, TD adolescents *n* = 33, and TD adults *n* = 60. Between-group comparisons of distance, HR intensity, and Borg RPE were performed using one-way ANOVA with Tukey’s post hoc test, or Kruskal-Wallis with Dunn’s test for non-normally distributed data. Differences in repeated-measures outcomes (D-F) over time and between groups were assessed using mixed-effects analysis; significant interactions were followed by post hoc comparisons using Dunnett’s test or Tukey’s test. $ indicates a significant time effect (difference from baseline) at a given time point, @ indicates a significant age effect (TD adolescents vs. TD adults) at a given time point, # indicates a significant difference between CP and TD adolescents at a given time point, and * indicates a significant difference between CP and TD adults at a given time point. Statistical significance was defined as adjusted p < 0.05. Data are presented as mean ± SD for each group unless otherwise indicated.

### Attenuated metabolic and oxygenation responses to acute exercise in adults with CP

The physiological effects of exercise were assessed by measuring venous blood gases at baseline, after 20 minutes of exercise, immediately post-exercise, and one-hour post-exercise. As expected, blood pH decreased modestly during exercise and returned toward baseline during recovery, with no clear differences between groups (Figure 3D). In contrast, several metabolic and gas exchange parameters differed with time and between groups during exercise and recovery (Figure 3D-F), indicating an overall attenuated systemic response in CP. More specifically, pCO_2_ and HCO_3_^-^ decreased less during exercise in CP, with differences observed relative to both TD groups at 20 minutes and relative to TD adults immediately post-exercise (Figure 3D). Individuals with CP also exhibited lower venous oxygen saturation and venous partial pressure of oxygen during and immediately after exercise compared with both TD groups, indicating differences in venous oxygenation throughout the exercise bout (Figure 3E). Lactate increased in all groups during exercise (mean post-exercise levels 5.1 ± 3.6 vs 5.3 ± 2.4 vs 7.5 ± 2.9 mmol/L in CP, TD adolescents, and TD adults, respectively) but did not differ between CP and either TD group at any individual time point (Figure 3F). Hemoglobin concentration and hematocrit increased transiently during exercise in all groups (Figure S5).

### Adults with CP exhibit an altered plasma metabolite response to acute exercise

Based on previous time-course studies identifying one hour as a responsive post-exercise timepoint (50), plasma metabolomic profiling identified exercise-associated changes in circulating metabolite levels across all groups compared to baseline, although the pattern of altered metabolites varied between groups (Figure 4A). Differential responses were observed in metabolites associated with TCA cycle metabolism, including lactate, malate, and α-ketoglutarate (α-KG). These metabolites showed a coordinated recovery response in TD participants, whereas this response was attenuated in individuals with CP, resulting in lower post-exercise levels of malate and α-KG in CP adults compared with TD adults (Figure 4B). Notably, the TCA-derived immunometabolite, itaconic acid (itaconate), exhibited a marked increase during recovery in adults with CP, whereas little or no change was observed in the TD groups (Figure 4B). Additional metabolites exhibiting group-dependent recovery responses are presented in Figure S6A-B.

**Figure 4.**
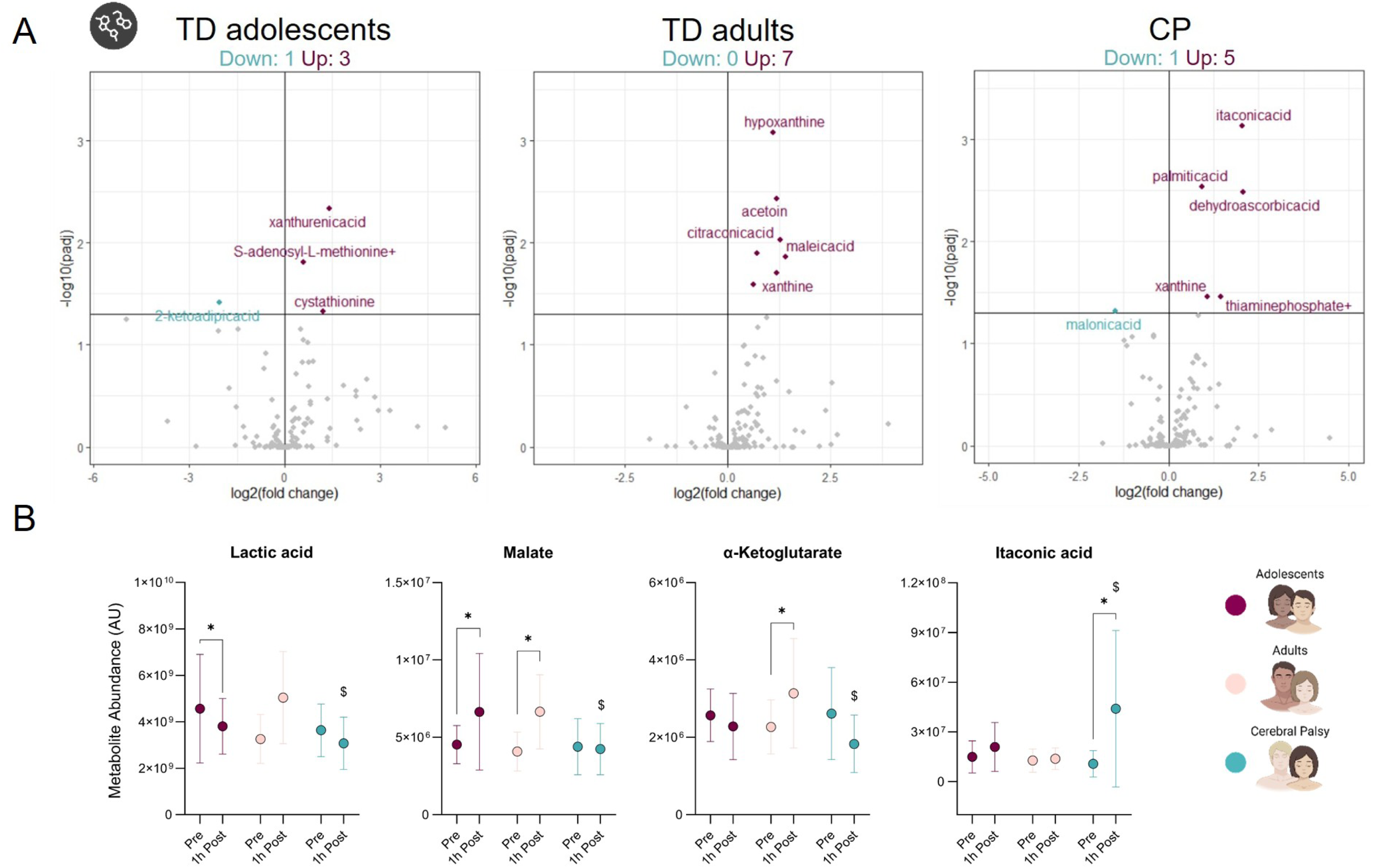
Plasma metabolite changes in CP and TD individuals in response to an acute bout of endurance exercise. (A) Volcano plots of all measured metabolites. The x-axis shows log2 fold change (FC), and the y-axis shows −log10 adjusted p values (Benjamini-Hochberg correction). Increased metabolites are shown in purple and decreased metabolites in green. The number of significantly increased and decreased metabolites is indicated above each volcano plot and the five most differentially abundant metabolites are annotated. (B) Changes in selected metabolites related to energy metabolism and the TCA cycle before and after exercise in TD adolescents, TD adults, and CP adults. Sample size: CP *n* = 12, TD adolescents *n* = 10, and TD adults *n* = 13. Statistical significance of changes in metabolite levels in response to exercise was assessed using two-way ANOVA. P values were adjusted for multiple comparisons using Tukey’s post-hoc test. * indicates a significant pre- to post-exercise difference, $ indicates a significant difference between CP adults and TD adults at post-exercise. Statistical significance was defined as adjusted p < 0.05. Data are presented as mean ± SD for each group.

### Individuals with CP show an attenuated plasma proteome response to acute exercise

To determine whether the altered physiological response to exercise in individuals with CP was reflected at the systemic level, we applied the same targeted plasma protein panel to samples one-hour post-exercise. This timepoint was selected as circulating exercise-responsive proteins exhibit substantial changes at this stage of the recovery process (51, 52). Overall, a robust exercise-induced response was observed in TD adolescents and adults, with 36 and 51 proteins, respectively, showing increased abundance at this timepoint. In contrast, individuals with CP exhibited a markedly blunted response, with 12 proteins showing increased abundance and one protein showing decreased abundance (Figure 5A).

**Figure 5.**
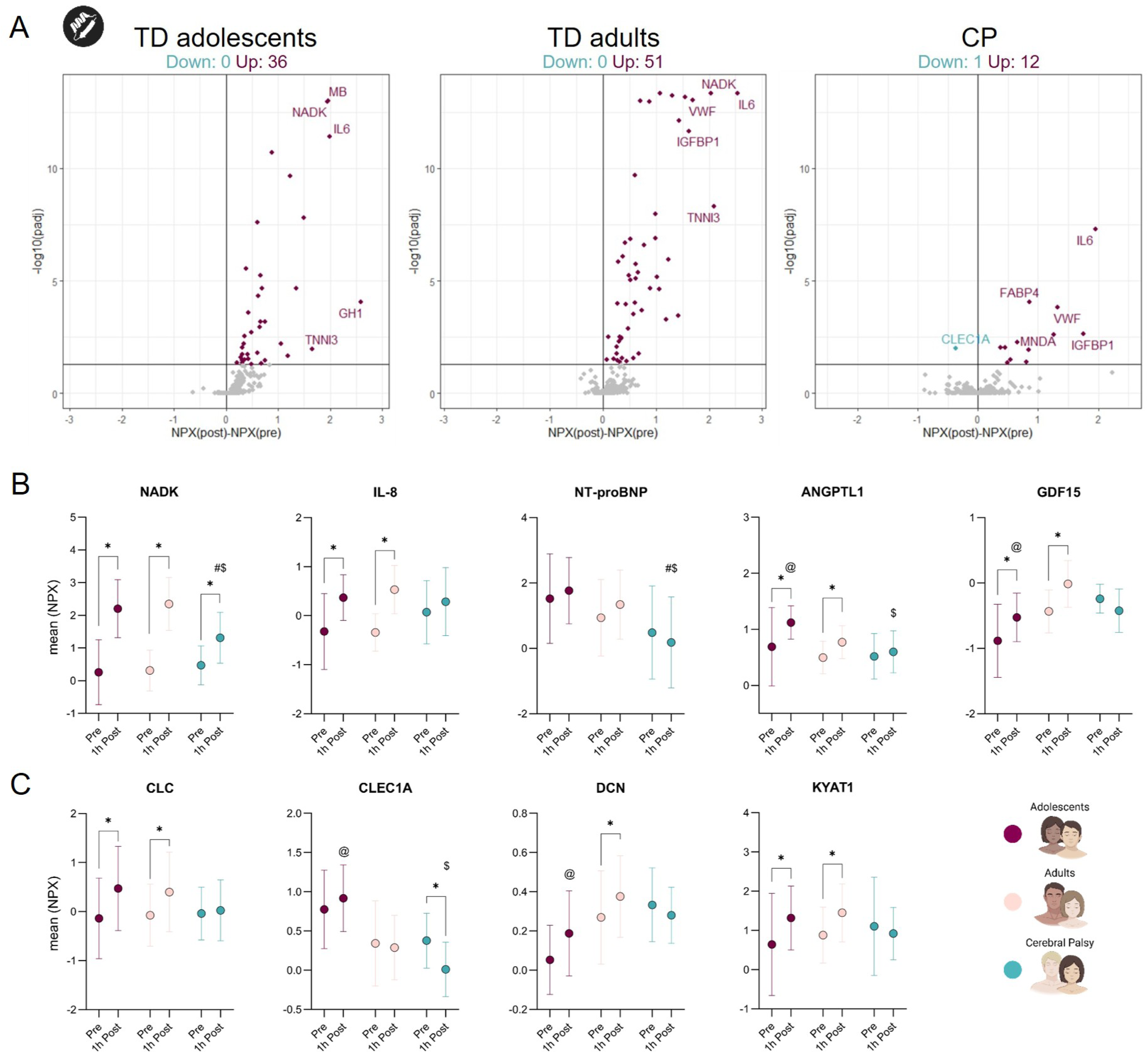
Plasma protein changes in CP and TD individuals in response to an acute bout of endurance exercise. (A) Volcano plots of all measured proteins. The x-axis shows delta NPX values and the y-axis shows −log10 adjusted p values (Benjamini-Hochberg correction). Increased proteins are shown in purple and decreased proteins in green. The number of significantly increased and decreased proteins is indicated above each volcano plot and the five most differentially abundant proteins are annotated. (B) Changes in selected exercise-responsive proteins before and after exercise in TD adolescents, TD adults, and CP adults. (C) Changes in selected proteins related to inflammation, the immune system, and ECM remodeling before and after exercise in TD adolescents, TD adults, and CP adults. Sample size: CP *n* = 15, TD adolescents *n* = 30, and TD adults *n* = 54. Statistical significance of changes in protein levels in response to exercise was assessed using two-way ANOVA. P values were adjusted for multiple comparisons using Tukey’s post-hoc test. * indicates a significant pre- to post-exercise difference, $ indicates a significant difference between CP adults and TD adults at post-exercise, # indicates a significant difference between CP adults and TD adolescents at post-exercise, @ indicates a significant difference between TD adolescents and TD adults at post-exercise. Statistical significance was defined as adjusted p < 0.05. Data are presented as mean ± SD for each group.

Individual proteins of biological relevance were then examined to further explore these systemic response patterns, including markers related to cardiometabolic function, vascular biology, immune regulation, and tissue remodeling. NAD kinase (NADK) and IL-8, both exercise-responsive factors (53, 54), increased following exercise in TD participants, whereas these responses were attenuated in CP. Similarly, N-terminal pro-B-type natriuretic peptide (NT-proBNP) and angiopoietin-like protein 1 (ANGPTL1), markers of cardiac stress and vascular remodeling (55), increased following exercise in TD participants, but not in those with CP, resulting in higher post-exercise levels in TD groups. Growth differentiation factor 15 (GDF15), another stress-responsive cytokine (56), also showed a marked post-exercise increase in TD participants but not in CP, consistent with an attenuated stress response (Figure 5B). Given basal alterations in immune-related proteins, we examined whether post-exercise immune signaling responses were altered in CP. Plasma levels of the immune-related protein CLC (galectin-10) increased following exercise in TD participants but not in CP. In contrast, plasma levels of the immune-related protein CLEC1A decreased following exercise in CP, resulting in lower post-exercise levels than those observed in TD adults (Figure 5C). This pattern was further supported by additional immune-related proteins, including SIGLEC7, CEBPB, and LGALS1, which displayed reduced or altered exercise responsiveness in CP compared to reference groups (Figure S7A-C). Beyond immune signaling, exercise also revealed group-specific regulation of proteins involved in ECM remodeling. Decorin (DCN), together with other regulators of ECM organization, displayed distinct exercise-induced responses between groups (Figure 5C, Figure S7A-C). Similar differences were observed across metabolic pathways. For example, KYAT1, a key enzyme in tryptophan-kynurenine metabolism, increased following exercise in TD participants but not in CP, consistent with differential regulation of circulating metabolites involved in tryptophan degradation (Figure 5C). Proteins involved in carbohydrate and lipid metabolism also exhibited distinct post-exercise patterns in CP (Figure S7A-C).

### The attenuated exercise response in CP persists when relative exercise intensity is matched

Lastly, we assessed whether differences in the acute exercise response persisted when relative exercise intensity was matched between groups. To achieve this, participants were selected based on heart rate-derived exercise intensity to obtain comparable relative intensities across groups. Heart rate intensity and perceived exertion did not differ between groups, yet adults with CP still covered a shorter distance than both TD groups (Figure 6A). Blood gas parameters changed over time in all groups (Figure 6B). pCO_2_ was higher in adults with CP than in TD adolescents at 20 minutes and higher than in TD adults immediately post-exercise. Venous pO_2_ was lower in CP than in both TD groups at 20 minutes and lower than in TD adults immediately post-exercise. Lactate increased during exercise in all groups and did not differ between CP and either TD group, although post-exercise lactate was higher in TD adults than in TD adolescents. At the circulating protein level, previously observed exercise-responsive patterns were largely maintained after intensity matching (Figure 6C). Myoglobin (MB) and NADK increased in all three groups, but post-exercise levels remained lower in adults with CP than in both TD groups. IL-8 increased in both TD groups but not in CP, while the patterns of the remaining selected proteins were largely preserved. These findings suggest that the blunted systemic and molecular response to exercise in CP was largely maintained at a comparable relative intensity.

**Figure 6.**
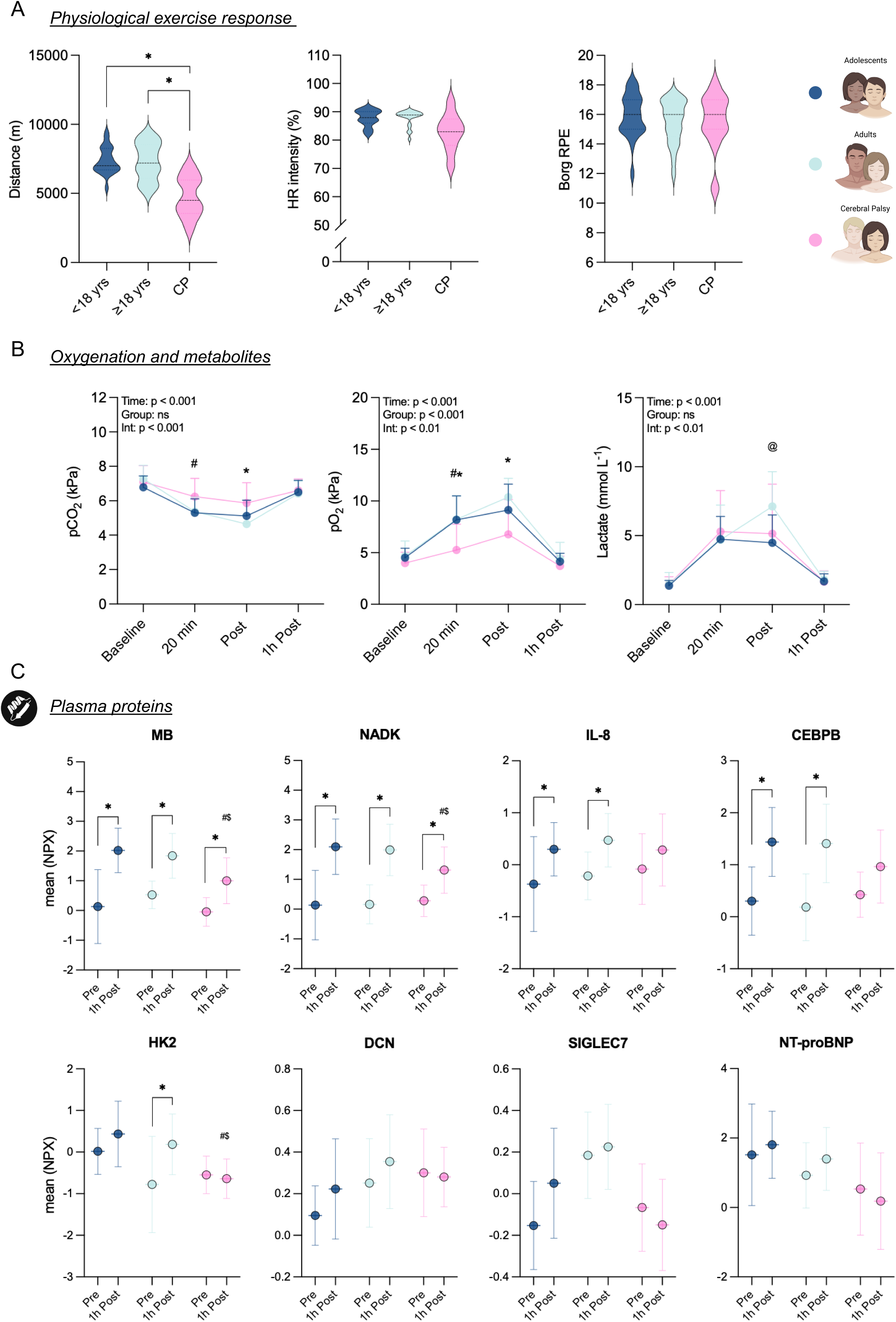
Physiological, metabolic, and plasma protein changes in CP and TD individuals in response to an acute bout of endurance exercise at matched relative exercise intensity. (A) Distance covered, mean heart rate intensity (% of maximal heart rate), and weighted mean Borg RPE during the session. Violin plots display the distribution of individual values, with the median indicated by a dashed line. (B) Partial pressure of carbon dioxide (pCO_2_), partial pressure of oxygen (pO_2_), and lactate at baseline, at 20 minutes, immediately post-exercise, and one-hour post-exercise. (C) Changes in selected exercise-responsive proteins before and after exercise. Sample size: CP n = 12, TD adolescents n = 21, and TD adults n = 21. Between-group comparisons of distance, HR intensity, and Borg RPE (A) were performed using one-way ANOVA with Tukey’s post hoc test, or Kruskal-Wallis with Dunn’s test for non-normally distributed data. Differences in repeated-measures outcomes (B) over time and between groups were assessed using mixed-effects analysis; significant interactions were followed by post hoc comparisons using Dunnett’s test or Tukey’s test. @ indicates a significant age effect (TD adolescents vs. TD adults) at a given time point, # indicates a significant difference between CP and TD adolescents at a given time point, and * indicates a significant difference between CP and TD adults at a given time point. Statistical significance of changes in protein levels (C) in response to exercise was assessed using two-way ANOVA. P values were adjusted for multiple comparisons using Tukey’s post-hoc test. * indicates a significant pre- to post-exercise difference, $ indicates a significant difference between CP adults and TD adults at post-exercise, # indicates a significant difference between CP adults and TD adolescents at post-exercise. Statistical significance was defined as adjusted p < 0.05. Data are presented as mean ± SD for each group unless otherwise indicated.

## DISCUSSION

Adults with CP experience a disproportionately high burden of secondary health complications, yet the underlying physiological mechanisms remain poorly understood. This knowledge gap is partly attributable to the longstanding view of CP as a pediatric condition, which has contributed to the underrepresentation of adults in clinical research (9). To address this gap, we integrated clinical, physiological, and molecular analyses to characterize the cardiometabolic physiology of adults with CP. Our findings identify impairments spanning multiple systems, including cardiovascular structure, systemic inflammation, metabolism, and the response to exercise. More specifically, adults with CP exhibited markedly reduced aerobic capacity together with structurally smaller heart dimensions. These physiological differences were accompanied by systemic molecular changes, exhibiting a circulating protein signature indicative of inflammation, ECM remodeling, and altered abundance of proteins associated with muscle biology. Furthermore, whereas a single bout of endurance exercise elicited robust physiological and molecular responses in TD adults, these responses were consistently blunted in adults with CP. These findings provide novel insight into the physiology of adults with CP, with potential clinical relevance for understanding cardiometabolic risk and impaired exercise responsiveness in this population.

To our knowledge, our findings provide the first characterization of cardiac structure in adults with CP and reveal smaller left and right ventricular dimensions compared with TD adolescents and adults. This extends previous work in younger cohorts, where structural differences have not been consistently observed (57), and suggests that differences in cardiac size may become more apparent in adulthood. However, most differences were attenuated after accounting for body size, indicating that the smaller cardiac dimensions observed in adults with CP may be due to smaller body sizes (58, 59). Despite these structural differences, resting systolic and diastolic function was still observed within expected ranges. Notably, left ventricular end-diastolic diameter was positively associated with VO_2_ peak in adults with CP, consistent with associations between ventricular dimensions and aerobic capacity in individuals without CP (60, 61). This suggests that the link between cardiac structure and aerobic capacity is preserved in adults with CP despite neurological impairment.

The mechanistic basis of these cardiac structural changes remains uncertain. Chronically reduced hemodynamic loading is associated with smaller ventricular chambers despite preserved systolic function (61–63), whereas endurance training produces the opposite response through sustained increases in preload (64–66). By analogy, chronically reduced cardiovascular stimulation due to lower physical activity may contribute to the smaller ventricular size observed in adults with CP (7, 67). Although habitual physical activity was not assessed, the lower circulating NPPB concentrations observed in adults with CP are consistent with a lower chronic cardiac loading state and provide indirect support for this interpretation. NPPB synthesis is primarily stimulated by myocardial wall stress and ventricular stretch (55), and recent studies further support a link between natriuretic peptides and ventricular morphology (68). Other CP-specific factors, including altered autonomic regulation and reduced muscle-pump activity, may also influence cardiac loading and filling (69–72).

Smaller ventricular dimensions would be expected to limit stroke volume and peak cardiac output (62, 63, 73). The reduction in RVDd, together with the blunted age-related increase in TAPSE, may also indicate reduced right ventricular reserve. Previous reports of reduced oxygen pulse in individuals with CP provide indirect support for impaired stroke volume augmentation during exercise (29, 74, 75). However, because most structural cardiac differences were no longer evident after indexing for body size and resting cardiac function remained preserved, the present findings do not support overt cardiac dysfunction as the primary explanation for reduced aerobic capacity. The proportionally greater reduction in skeletal muscle thickness in CP further suggests that reduced aerobic capacity may reflect both central and peripheral factors.

The venous blood gas response provides further insight into the exercise response in CP. During exercise, pCO_2_ and HCO_3_^-^ fell less in adults with CP. At higher exercise intensities, increased ventilation normally lowers pCO_2_ as part of respiratory compensation for metabolic acidosis (76). The smaller reductions in CP may therefore reflect a less pronounced systemic perturbation, consistent with the lower external workload and heart rate response. However, an altered ventilatory response may also contribute. Smaller tidal volumes accompanied by a higher respiratory frequency have previously been observed during exercise in individuals with CP (29), potentially increasing the relative contribution of dead-space ventilation and attenuating the fall in pCO_2_. In contrast, venous oxygen saturation and pO_2_ were lower in CP during and immediately after exercise, despite the lower external workload. As venous blood reflects metabolic events in the tissue it drains (76), this may indicate an altered interaction between oxygen demand, blood flow, and oxygen extraction. This is consistent with the lower oxygen pulse observed in CP, which may reflect differences in stroke volume and/or peripheral oxygen extraction (29). Differences in oxygen delivery, fractional extraction, and muscle recruitment during Frame Running may contribute to the lower venous oxygen values. Our findings thus suggest altered coordination of cardiovascular, metabolic, and peripheral oxygen responses in CP.

Compared to TD adults, those with CP showed signs of a pro-inflammatory circulating profile, consistent with the concept of low-grade systemic inflammation (77, 78). Notably, these findings were observed in a cohort of adults with CP who were of normal weight, suggesting that the inflammatory profile was not simply associated with overweight/obesity. This is important when considering the broader CP population, in which greater adiposity may further contribute to inflammatory burden (79). When compared with the TD adolescent group, however, the differences were less pronounced, with some markers in TD adolescents being comparable to those observed in adults with CP. This may reflect age- and puberty-related differences in immune regulation, as circulating inflammatory cytokines can vary across childhood, adolescence, and early adulthood (43, 44). Several factors could contribute to systemic inflammation in CP, including physical inactivity and other consequences of living with the condition. However, the limited overlap between the basal plasma protein profile and proteins associated with habitual physical activity provides little support for inactivity as a major driver. Recent findings from our group indicate that systemic inflammation is not consistently present in young children with CP (manuscript under review), suggesting that it may emerge or become more pronounced with advancing age. If sustained, such inflammation may contribute to the increased chronic disease burden observed in this population, with potential effects on both cardiometabolic (80, 81) and skeletal muscle health (82).

Beyond the cardiovascular phenotype, adults with CP exhibited reduced skeletal muscle thickness compared with TD controls, consistent with previous reports of impaired muscle development and reduced muscle mass in CP (83, 84). Given its role in whole-body metabolic regulation (26), reduced muscle mass may contribute to the elevated cardiometabolic risk observed in adults with CP (77, 85), and limit aerobic capacity by reducing the tissue available for oxygen utilization during exercise (24, 86). Lower muscle mass may reflect altered molecular regulation of skeletal muscle. For instance, relative to TD adults, those with CP had elevated circulating myostatin, a potent negative regulator of muscle mass (87, 88). This finding extends earlier reports of increased skeletal muscle myostatin expression in children with CP (89), by providing evidence that this pathway may also be altered at the systemic level. However, myostatin levels did not differ from the TD adolescent group, potentially reflecting a developmental increase associated with puberty (37). Additionally, circulating DLK1, a recently described myostatin antagonist that limits Activin receptor-mediated pathway activation (42), was markedly reduced in the CP group compared with both TD reference groups, providing further evidence of altered regulation of this pathway. Other circulating factors, such as ECM proteins and inflammatory markers, may also contribute to the smaller muscle size observed in CP, mirroring mechanisms described in aging and clinical cohorts (82, 90–92).

Our findings reveal systemic metabolic differences in adults with CP, evident both at rest and after exercise. While alterations in circulating metabolites have been reported in pediatric CP cohorts (93, 94), the specific metabolites and their direction of change differ from those observed here, indicating that the CP metabolic phenotype is not static and may shift across the lifespan. In the present study, adults with CP had lower resting circulating levels of aconitic acid and 2-HG compared with both TD groups. Given their close ties to the TCA cycle, the reduced abundance of these molecules may reflect alterations in mitochondrial bioenergetics, supported by evidence of compromised mitochondrial function in CP skeletal muscle (32, 95–97). Moreover, in response to physical exercise, the CP group showed blunted or absent changes in lactate, malate, and α-ketoglutarate compared with TD controls. These data provide evidence of altered exercise metabolism in this population under standardized conditions. Further examining these data, we also observed a marked post-exercise rise in itaconic acid (itaconate), a metabolite with immunoregulatory and metabolic functions (98, 99). Synthesized from cis-aconitate via ACOD1/IRG1, prior work has shown that itaconate serves as an anti-inflammatory “brake” that suppresses excessive macrophage activation during inflammatory stress (99). Given the elevated inflammatory baseline in CP, the marked post-exercise increase in itaconate may thus represent a counter-regulatory response to exercise-induced inflammatory stress.

The generally attenuated molecular response to exercise in adults with CP might reflect several, potentially interacting, mechanisms. First, differences in the basal state may constrain the magnitude of the subsequent response. Given that several proteins and metabolites were already altered at baseline in CP, these differences may restrict further responses to acute physiological stress. Such a “ceiling effect” has been reported in other conditions (e.g., myalgic encephalomyelitis/chronic fatigue syndrome) with altered molecular profiles (100, 101). Second, differences in tissue composition and metabolic activity may shape the circulating response. Exercise-responsive factors, including exerkines (102), as well as metabolites released or consumed by skeletal muscle, adipose tissue, liver, and immune cells, may differ according to tissue mass, composition, and metabolic activity. Finally, differences in the exercise stimulus could potentially contribute to the attenuated molecular response. To address this, we repeated the analyses after restricting the TD groups to participants who achieved comparable heart rate responses to the CP group. Most of the observed physiological and molecular patterns remained, suggesting that differences in exercise intensity do not primarily explain the attenuated response. However, differences in the exercise stimulus cannot be fully excluded, given the distinct biomechanical demands of Frame Running compared with conventional running.

Structured exercise training is an important strategy to improve cardiometabolic health and reduce secondary complications in individuals with CP (3, 30). Beneficial effects of exercise training have been shown in this population across multiple physiological domains, including improvements in aerobic capacity, muscle function, and cardiovascular health (103–105). However, most of these studies have not included a TD control group, and it therefore remains unclear whether the magnitude and nature of training adaptations in adults with CP are comparable to those observed in TD individuals. Our findings of altered physiological and molecular responses to acute exercise raise important questions about whether, and to what extent, these differences may translate into altered adaptations to long-term training and, ultimately, long-term health. Future research should therefore determine whether exercise strategies developed in the general population can be directly applied to adults with CP or whether they require further tailoring to account for different exercise responses, biomechanical inefficiencies, and trainability. Such work will be important for refining exercise prescription and ensuring that training elicits clinically meaningful health benefits in this population.

Some limitations should be considered when interpreting these findings. The cross-sectional design precludes causal inference and limits our ability to determine whether the observed differences reflect the condition itself, underlying differences in habitual physical activity, neuromuscular factors such as agonist-antagonist coactivation, rate coding, and motor unit recruitment, or a combination of these factors. The CP cohort was also heterogeneous with respect to GMFCS level and subtype, which may mask subgroup-specific effects that could be identified in larger, more stratified cohorts. The use of different exercise modalities is another limitation. Participants with CP used running frames, whereas TD participants performed conventional running, introducing potentially different biomechanical and neuromuscular demands. Although we repeated the analyses using TD participants with comparable heart rate responses, which recapitulated the group-specific patterns, the effects of exercise modality and external workload cannot be fully separated from those of group. Interpretation of the cardiac findings also requires consideration of body size, as most structural differences were attenuated after indexing to body surface area. In addition, variation in echocardiographic image quality may have affected the accuracy of cardiac dimensional measurements. Cardiac magnetic resonance imaging would provide a more robust assessment in future studies. The cross-sectional design also means that we cannot determine how cardiac structure develops across the lifespan or whether the observed characteristics reflect early differences, long-term (mal)adaptations, or age-related changes. Finally, the molecular responses were assessed at predefined post-exercise time points. Given the time-dependent nature of proteomic and metabolomic responses, different sampling strategies may capture additional or different changes depending on the markers examined.

In conclusion, this study demonstrates that adults with CP exhibit a physiological profile that is distinct from that of TD individuals, both at baseline and in response to an acute exercise challenge. At baseline, this profile is characterized by reduced aerobic capacity together with differences in cardiac structure, skeletal muscle mass, and the systemic milieu, suggesting diminished physiological reserve across multiple biological systems. Following acute exercise, adults with CP also exhibit consistently attenuated physiological and molecular responses, indicating an altered capacity to respond to exercise-induced stress. Whether these differences reflect CP-related biological mechanisms, long-term differences in physical activity and conditioning, differences in the exercise stimulus, or a combination of these factors remains to be determined. Together, these findings provide novel insight into the biological basis of cardiometabolic vulnerability in adults with CP and suggest that conventional exercise strategies may require further refinement to optimize health and fitness in this population.

## METHODS

### Sex as a biological variable

The study included both males and females. Due to the limited sample size in the CP cohort, sex was not included as a biological variable in the analyses. However, sex was evenly distributed across the three groups to reduce the potential for sex-related confounding.

### Study design and participants

This observational study employed a cross-sectional design with integrated cardiometabolic phenotyping in individuals with cerebral palsy (CP) and typically developing (TD) adolescents and adults. Data were collected across several study phases between 2018 and 2024, involving overlapping but not identical participant subsets for different assessments. Across one or more study components, a total of 117 individuals were included, contributing 121 observations: 23 individuals with CP (27 observations, aged 18-44 years), 33 TD adolescents (aged 13-17 years), and 61 TD adults (aged 18-50 years). Individuals with CP were included if they had a confirmed diagnosis of CP, were able to understand and follow verbal instructions, and had sufficient experience to exercise continuously using a running frame for at least 10 minutes. TD participants self-identified as healthy and reported no regular use of prescribed medication. Exclusion criteria applied to all participants included recent botulinum toxin injections (≤3 months), soft tissue surgery (≤6 months), bone surgery (≤12 months), and pain or medical conditions likely to affect physical performance or study participation. Participants with CP were recruited during World Abilitysport International Frame Running Camp and Cup events in Copenhagen in 2018 and 2019, with additional recruitment conducted in Sweden in 2022 through Frame Running clubs, supervised training sessions, and social media outreach. Some individuals with CP took part on more than one occasion. Two individuals were sampled twice, four years apart, and one individual was sampled three times over a four-year period, resulting in 27 observations from 23 individuals with CP. Each sampling occasion was treated as a separate observation in the analyses. Not all participants completed all assessments; therefore, sample sizes vary by outcome and are specified in Table S1. An overview of participant characteristics is provided in Table 1.

### Anthropometric measurements

Body weight was measured using a wheelchair-accessible electronic scale (Detecto 6550, Detecto Scale Company, Web City, MO, USA). Body height was assessed using a flexible tape measure, with participants positioned either standing or supine on an examination table depending on functional ability. In participants with joint contractures or limited range of motion preventing full extension, segmental measurements between anatomical landmarks were summed to estimate total body height. Body weight and height were measured using the same procedures described above in all TD participants and in all CP observations, except for 14 CP observations in which body height and weight were self-reported. These data are summarized in Table 1.

### Cardiac assessments

Resting cardiopulmonary function was assessed using a standard 12-lead electrocardiogram (ECG) (IntelliVue MX550 Patient Monitor, Philips Healthcare, Amsterdam, the Netherlands) and transthoracic echocardiography (Vivid S70, GE Healthcare, Chicago, IL, USA). All recordings were obtained in the supine position following established clinical guidelines. Resting blood pressure was measured after a period of supine rest. Resting ECG was used to evaluate heart rate, PQ interval, QRS duration and corrected QT interval (QTc). Transthoracic echocardiography was performed to assess cardiac structure and function, including global longitudinal strain (GLS), ejection fraction (EF) using Simpson’s biplane method, interventricular septal thickness at end-diastole (IVSd), left ventricular diameter at end-diastole (LVDd), right ventricular diameter at end-diastole (RVDd), tricuspid annular plane systolic excursion (TAPSE), left atrial volume index (LAVi), systolic myocardial velocity (S’), and indices of left ventricular diastolic function and filling pressures (E/e’ and E/A ratio). To account for differences in body size between groups, LVDd, RVDd, and IVSd were indexed to body surface area (BSA) and expressed as LVDi, RVDi, and IVSi (mm/m^2^). BSA was calculated from body height and body mass using the Haycock formula (106). All examinations were conducted by experienced clinicians and interpreted according to standard clinical criteria.

### Skeletal muscle thickness assessment

Skeletal muscle thickness of the *m. rectus femoris* was assessed using B-mode ultrasonography, following an established protocol (19). Measurements were performed using an ultrasound system (EPIQ 7, Philips Healthcare, Bothell, WA, USA) equipped with a linear array transducer. Images were analyzed using MicroDicom Viewer DICOM (64-bit), where the antero-posterior distance between the superficial and deep aponeuroses of the *m. rectus femoris* was measured without probe-induced compression of the skin or underlying muscle. Two images were analyzed, and the mean of the two measurements was used for analysis. Muscle thickness data for a subset of participants (CP n=13 and TD n=30) have been reported previously (19).

### 6-Minute Frame Running Test

Participants with CP performed the 6-Minute Frame Running Test (6-MFRT) (107, 108), a field-based test in which participants use a running frame to cover the maximum possible distance in 6 minutes, as previously described (107–110). Heart rate, Borg Rating of Perceived Exertion (RPE), and distance covered were recorded throughout the test, with distance used as a validated proxy for CRF (109). Of the 27 CP observations, 22 included a valid test. In one observation, the criteria for a valid maximal physiological effort were not met, and in four the test was not performed or data were missing.

### Peak oxygen uptake test

Peak oxygen uptake (VO_2_ peak) was assessed using an incremental exercise test performed to volitional exhaustion. Participants with CP performed the test using a running frame on a wide treadmill (Rodby v2, Rodby Innovation AB, Vänge, Sweden), whereas TD participants performed conventional treadmill running (Rodby RL 2000E or 2500E, Rodby Innovation AB, Sweden). Pulmonary gas exchange was measured continuously on a breath-by-breath basis throughout the test using a calibrated metabolic cart (Quark CPET, COSMED, Rome, Italy). Heart rate was recorded continuously using a chest-strap heart rate monitor (SmartLAB HRM, Heddesheim, Germany). Individualized exercise protocols were applied for participants with CP to accommodate differences in motor function and endurance capacity. Of the 23 adults with CP, 12 completed the maximal exercise test, of whom 10 fulfilled the criteria for a valid maximal effort. Of the remaining 11 participants, 10 were recruited during the World Abilitysport International Frame Running Camp and Cup in Copenhagen, where maximal exercise testing was not part of the study protocol, and one participant recruited in Sweden did not perform the test. Treadmill speed and/or inclination were increased incrementally until volitional exhaustion. TD participants performed a standardized incremental treadmill protocol to task failure (i.e., until voluntary cessation). Strong verbal encouragement was provided throughout all tests. Criteria for achieving maximal effort, data processing procedures, and protocol details have been described elsewhere (29). VO_2_ peak was defined as the highest 30-second rolling mean value obtained during the test and is reported as an absolute value (L/min) and relative to body mass (mL/kg/min).

### 45-minute endurance exercise session

Individuals with CP and TD participants completed a 45-minute endurance exercise session, as previously described (33, 111). The session was performed on a 200-m indoor track. Participants with CP used a running frame, whereas TD participants performed conventional running. Participants were instructed to cover the maximum possible distance while maintaining a perceived exertion corresponding to ≥15 on the Borg RPE scale throughout the session. Borg RPE was assessed at 10, 20, 30, 40, and 45 minutes, and a weighted mean RPE across the session was subsequently calculated for each participant. Heart rate and distance covered were monitored continuously using a chest-strap heart rate monitor (Garmin, Olathe, KS, USA). Heart rate values were recorded as one-minute averages at 10-minute intervals and immediately upon completion of the session. A weighted mean heart rate was calculated and expressed relative to maximal heart rate. As the 45-minute endurance session was performed at a submaximal, self-paced intensity, maximal heart rate was defined as the highest value obtained during either the VO_2_ peak test or the 45-minute exercise session. If this value was lower than the age-predicted maximum (220 – age), the age-predicted formula was used instead. No standardized warm-up was performed prior to the exercise session. A subset of TD participants (n=64) consumed a standardized breakfast one hour prior to the exercise session, as previously described (111). Upon arrival at the exercise venue, a peripheral venous catheter was inserted into an antecubital vein in all participants. Participants with CP and TD adolescents applied local anesthetic cream (EMLA; Aspen Pharma Trading Limited, Dublin, Ireland) at least one hour before the session to minimize discomfort during blood sampling. Venous blood samples were collected at rest before exercise and one hour after completion of the 45-minute exercise session using EDTA collection tubes. Blood gas samples were obtained from venous blood at baseline, after 20 minutes of exercise, immediately after exercise, and one-hour post-exercise. Blood gas samples were analyzed immediately using a blood gas analyzer (ABL800 Flex®, Radiometer, Copenhagen, Denmark) to assess acid-base balance.

### Blood sample processing

Venous blood samples collected at rest and one-hour post-exercise were processed immediately after collection. Samples intended for proteomic and metabolomic analyses were centrifuged according to standard protocols, and plasma was aliquoted and stored at -80°C until analysis. All samples were thawed only once prior to downstream analyses.

### Plasma proteomics

Plasma protein profiling was performed using the antibody-based Olink Explore Cardiometabolic panel I (Olink Proteomics, Uppsala, Sweden) as it covers established biomarkers related to cardiovascular and metabolic health. As described previously (54), we quantified 366 individual proteins from this panel and reported their levels as normalized protein expression (NPX) values, Olink’s relative measure of protein abundance. Samples were processed as single measurements and analyzed in a randomized sequence. To reduce technical variability, pre- and post-exercise samples from the same participant were analyzed on the same plate, and normalization across plates was applied during data processing. Assay performance and analytical consistency were monitored using quality control samples included on each plate. Based on NPX values from pooled plasma controls, the intra- and inter-assay coefficients of variation were 14% and 16%, respectively.

### Plasma metabolomics

Plasma metabolomics was performed in line with procedures outlined in Hanaoka et al. 2025 (35). Briefly, total hydrophilic metabolites were extracted from plasma using an 80:20 methanol/water extraction protocol. After initial centrifugation, 200 µL of supernatant was mixed with 800 µL ice-cold methanol/water, vortexed, and centrifuged at 20,160 g for 15 minutes at 4°C. The extraction solution was dried by SpeedVac, reconstituted in 50% acetonitrile, and centrifuged prior to analysis. The resulting supernatant was used for targeted metabolomics of 261 core metabolites related to carbohydrate, lipid, and protein metabolism using high-performance liquid chromatography coupled with high-resolution tandem mass spectrometry.

### Statistical analysis

Descriptive data and clinical variables were assessed for normal distribution using a Shapiro-Wilk test. Between-group comparisons of participant characteristics, resting cardiopulmonary measures (blood pressure, ECG, and echocardiographic variables), and single-time-point exercise outcomes (distance, HR intensity, and Borg RPE) were performed at baseline using one-way analysis of variance (ANOVA) or Kruskal-Wallis tests, depending on data distribution. Baseline differences in Olink-derived plasma proteome and metabolome measures were also assessed using one-way ANOVA. When overall group effects were detected, post-hoc pairwise comparisons were conducted using Tukey’s test for parametric data or Dunn’s test with appropriate adjustment for multiple comparisons for non-parametric data. The association between VO_2_ peak, LVDd and rectus femoris thickness was assessed separately within each group using Pearson correlation coefficients (r).

Exercise-induced changes in acid-base balance and other repeated-measures outcomes obtained at multiple time points (pre-exercise, during exercise, immediately post-exercise, and one hour post-exercise) were analyzed using a two-way mixed ANOVA, or a mixed-effects analysis when data were not normally distributed, with time (within-subject) and group (CP vs TD adolescents vs TD adults, between-subject) as factors. Significant interactions (time or time vs group) were followed by post-hoc comparisons using Dunnett’s multiple comparison test, Tukey’s multiple comparisons test, or uncorrected Fisher’s LSD. For the Olink-derived plasma proteome and metabolome, acute exercise-induced changes were assessed using two-way ANOVA followed by Tukey post-hoc testing with adjustment for multiple comparisons. The level of statistical significance was set to *p* < 0.05. All statistical analyses were performed using GraphPad Prism (version 11.0.2 for Mac, GraphPad Software, Boston, MA, USA), and IBM SPSS Statistics (version 31.0.2.0, IBM Corp., Armonk, NY, USA). Plasma proteomic and metabolomic analyses were performed using R (version 4.5.1). A statistical summary is reported in Table S1.

### Study approval

The study was approved by the Swedish Ethical Review Authority (DNR 2016/1139-31/2, 2021-05116, 2023-02343-01), including approved addenda (DNR 2016/1675-32, 2017/2237-32, 2020-04170, 2022-06652-02, 2023-07692-02). All study procedures were conducted in accordance with the Declaration of Helsinki. All participants received oral and written information regarding the study procedures and the potential risks and benefits associated with participation. Written informed consent was obtained from all participants, or from legal guardians for participants younger than 18 years of age.

## Data availability

The data supporting the findings of this study will be deposited in Figshare upon submission.

## Acknowledgements

The authors sincerely thank all study participants. We also thank Riksidrottsförbundet (the Swedish Sports Confederation) for access to the testing facilities at Bosön, Christoffer Sundqvist for assistance with physiological testing, and Björn “Bloody-Bear” Otto Lanzky for assistance with blood sampling. Figures were created using BioRender, GraphPad, and R.

## ADDITIONAL INFORMATION

### Author contributions

FvW and JN conceived the study. LC, SE, SMR, WH, JC, AK, JS, EH, OK, PS, RFG, JN, and FvW recruited participants and collected samples. LC, OH, and SMR analyzed the data. LC and OH prepared the figures and wrote the original draft of the manuscript. DAC, PS, JN, and FvW provided resources and facilities. OH, SE, JN, and FvW acquired funding. All authors reviewed and edited the manuscript and approved the final version. LC and OH share first authorship and contributed equally to data analysis and manuscript preparation. LC is listed first owing to her role in participant recruitment and sample collection.

### Competing interest

The authors declare no competing interests.

### AI disclosure

Claude (Opus 5.5, Anthropic) and ChatGPT (OpenAI) were used to assist with data organization and with language editing during manuscript preparation (August-October 2026). The authors assume full responsibility for ensuring the integrity and accuracy of AI output presented in this article.

### Funding

OH was supported by grants from Svenska Sällskapet för Medicinsk forskning (PG-25-0562), Anna-Lisa och Arne Gustafssons Stiftelse, Stiftelsen Tornspiran, Folke Bernadotte Stiftelsen, Stiftelsen Sunnerdahls Handikappfond (F32/25), Stiftelsen Maja & J.P. Åhlén (253029), Stiftelsen Frimurare Barnhuset i Stockholm, Kronprinsessan Lovisas Förening för Barnasjukvård (2026–266), Sällskapet Barnavård (2026–077) and the Network for Movement & Physical Function. SE was supported by grants from Svenska Sällskapet för Medicinsk forskning (PG-24-0429), Linnea och Josef Carlssons Stiftelse, Stiftelsen Sunnerdahls Handikappfond, Sällskapet Barnavård, Stiftelsen Sven Jerrings Fond and KI Research Foundation Grants (#2024-02598). FvW was supported by the Swedish Research Council (no. 2022-01392), the Linnea and Josef Carlssons Foundation, the Norrbacka Eugenia Foundation, the ELSASS Foundation (24-B01 1022, 25-B01-1319), Neuroförbundet (SB2025-0005), and the Swedish Society for Medical Research (SG-25-0199). FvW and JN were supported by a strategic research grant from the Promobilia Foundation (S24502).

---

**Supplementary Figure 1.**
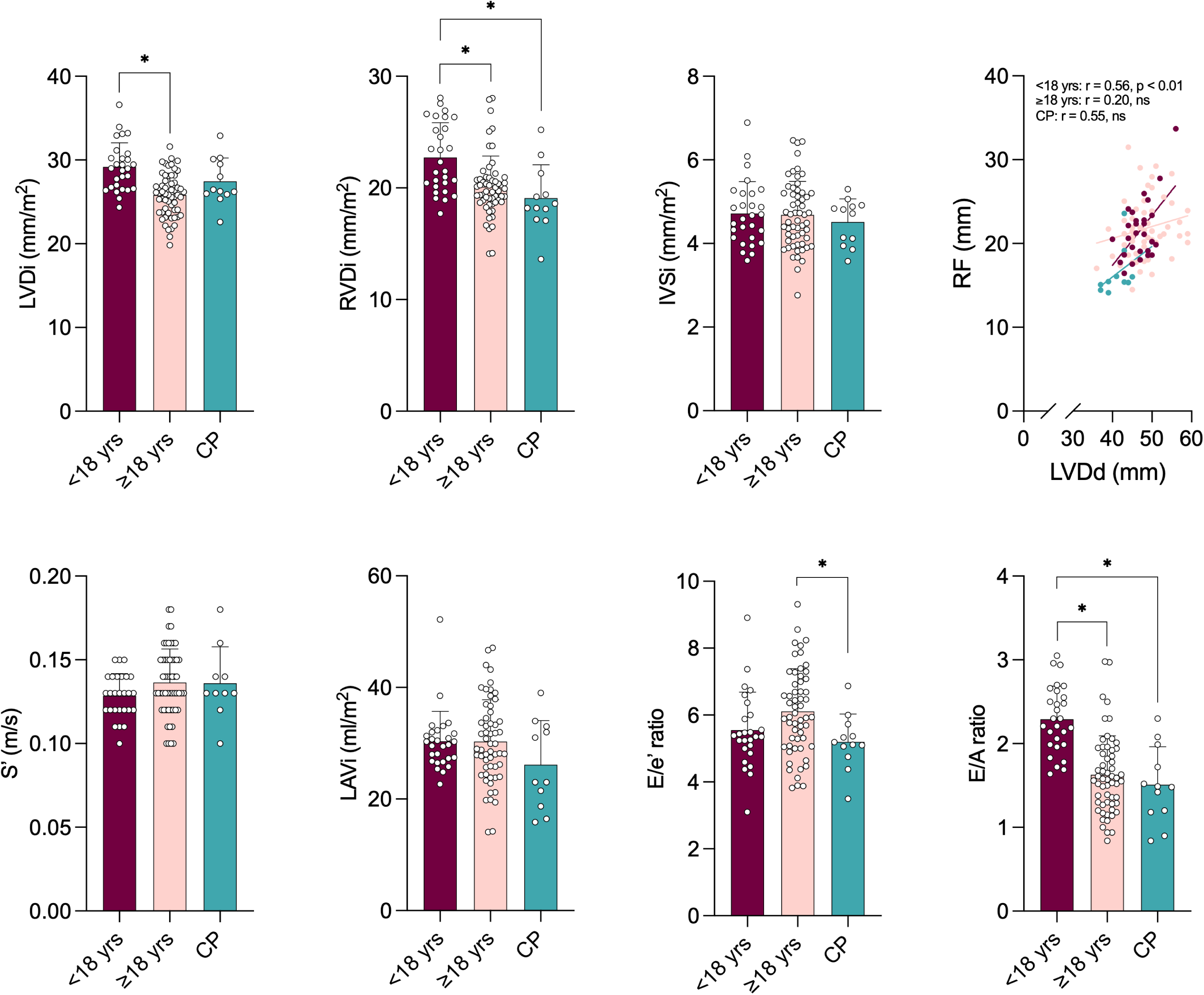
Cardiac dimensions indexed to body surface area, association between left ventricular size and muscle thickness, and additional systolic and diastolic indices in CP and TD individuals. Cardiac dimensions indexed to body surface area: left ventricular diameter at end-diastole index (LVDi), right ventricular diameter at end-diastole index (RVDi), and interventricular septal thickness at end-diastole index (IVSi). Correlation between left ventricular diameter at end-diastole (LVDd) and m. rectus femoris (RF) muscle thickness within each group, with Pearson correlation coefficients (r) and p values indicated. Systolic myocardial velocity (S’), left atrial volume index (LAVi), E/e’ ratio, and E/A ratio in TD adolescents, TD adults, and individuals with CP. Sample size: CP n = 12, TD adolescents n = 28, and TD adults n = 58. Significant differences between groups were assessed using one-way ANOVA with Tukey’s post hoc test, or Kruskal-Wallis with Dunn’s test for non-normally distributed data. * indicates adjusted p < 0.05. Data are presented as mean ± SD for each group.

**Supplementary Figure 2.**
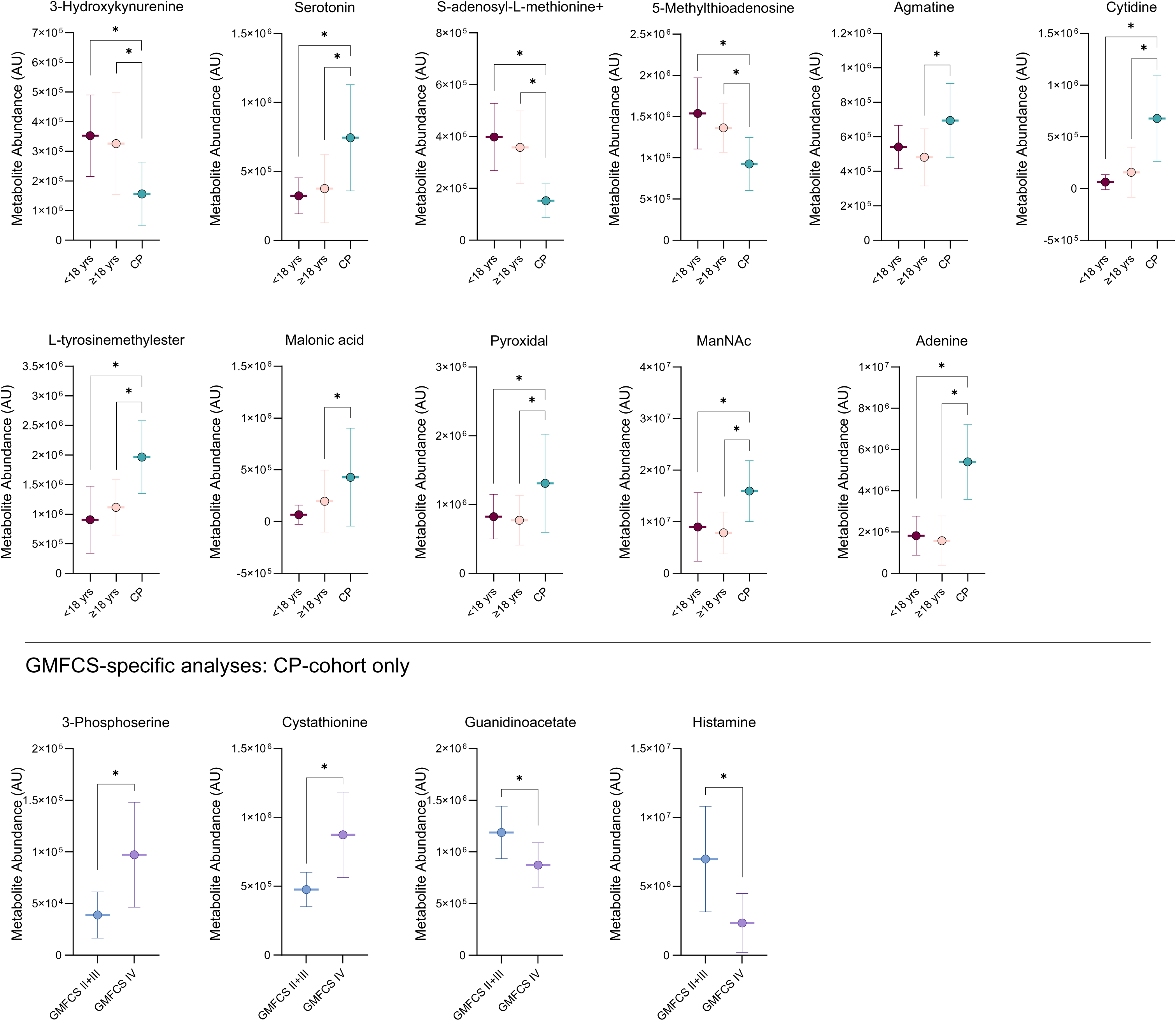
Additional basal circulating metabolite profiles in CP and TD individuals and GMFCS-specific metabolite analysis in CP. Sample size: CP *n* = 13, TD adolescents *n* = 13, and TD adults *n* = 17. Significant differences in metabolite levels between groups were assessed using one-way ANOVA. Differences in metabolite levels between GMFCS II+III and IV groups were assessed using unpaired t-tests. P values were adjusted for multiple comparisons using Tukey’s post-hoc test. * indicates adjusted p < 0.05. Data are presented as mean ± SD for each group.

**Supplementary Figure 3.**
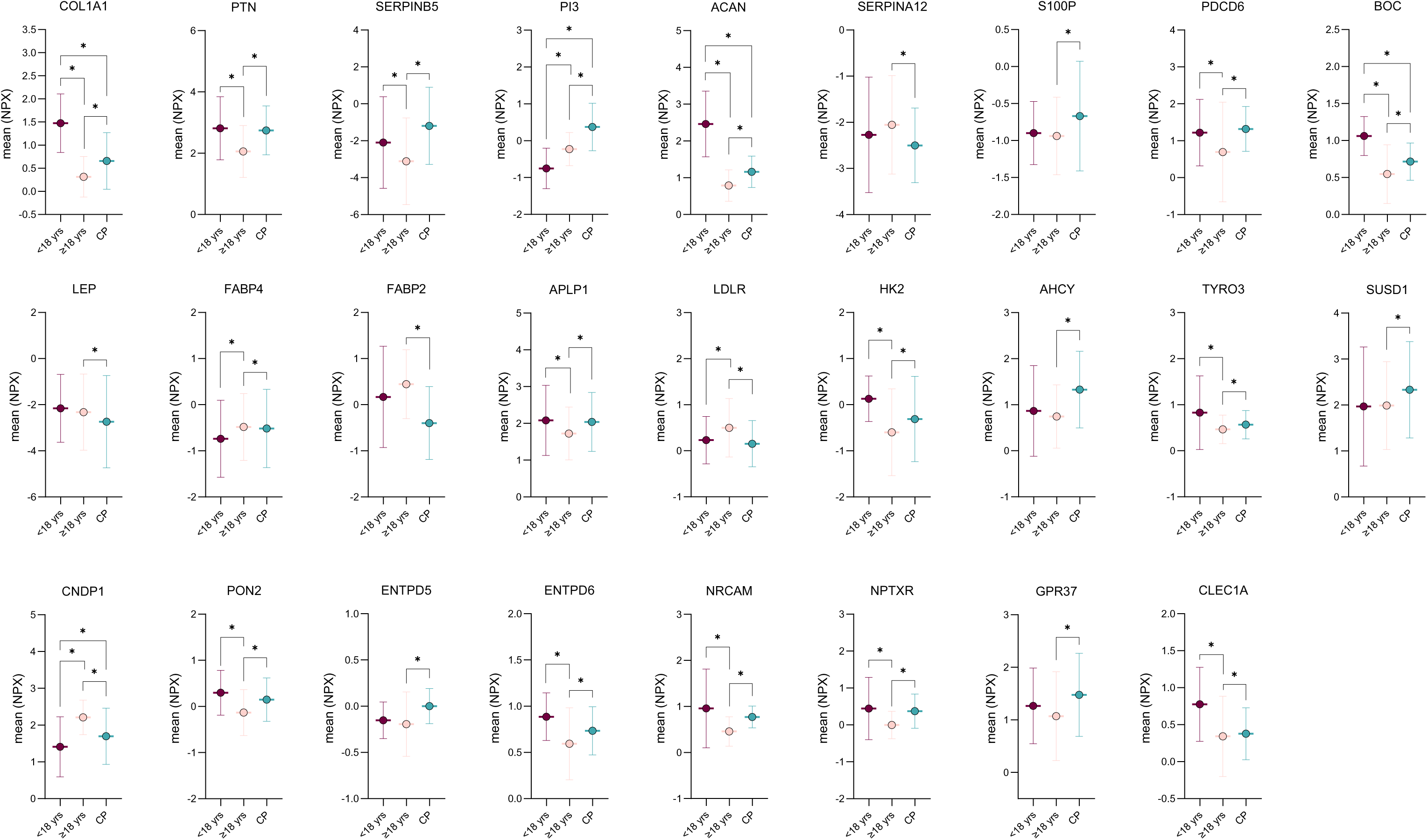
Additional basal circulating protein profiles in CP and TD individuals. Sample size: CP *n* = 27, TD adolescents *n* = 30, and TD adults *n* = 59. Significant differences in protein levels between groups were assessed using one-way ANOVA. P values were adjusted for multiple comparisons using Tukey’s post-hoc test. * indicates adjusted p < 0.05. Data are presented as mean ± SD for each group.

**Supplementary Figure 4.**
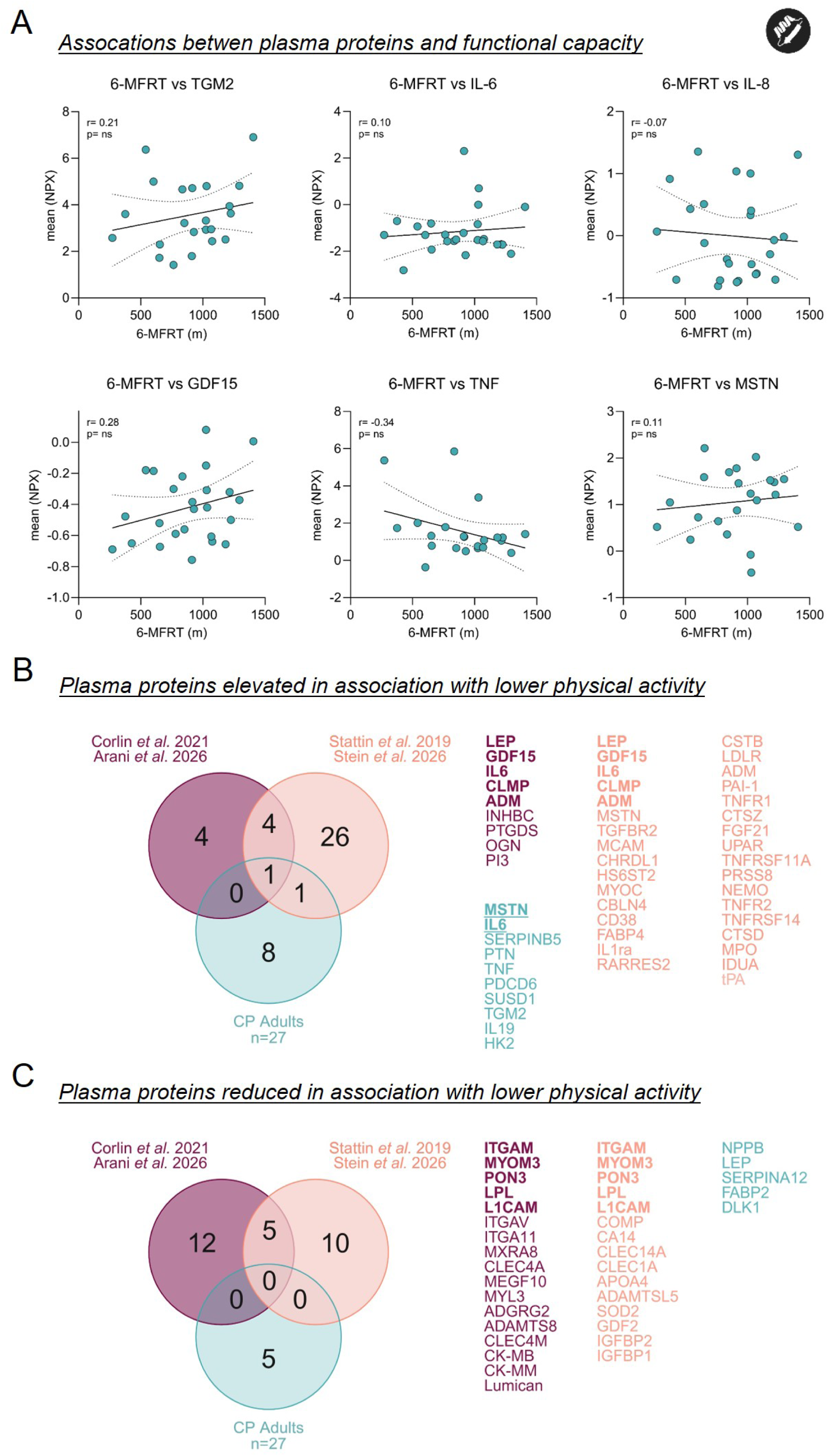
Relationship between plasma proteins, functional capacity, and physical activity-associated proteomic signatures. (A) Pearson correlations between 6-MFRT performance and selected plasma proteins (TGM2, IL-6, IL-8, GDF15, TNF, and MSTN) in adults with CP (n = 25). Pearson correlation coefficients (r) and P values are indicated. (B-C) Comparison of the 15 most differentially abundant plasma proteins in adults with CP relative to TD adults with proteins associated with physical activity in four published datasets (see references (45–48)), presented as Venn diagrams. The 15 CP-associated proteins comprised 10 increased and five decreased proteins. Proteins associated with physical activity were extracted from reported protein lists and figures, including proteins showing strong associations with physical activity and, where available, proteins replicated in independent cohorts within the respective studies. (B) Proteins elevated with lower physical activity. (C) Proteins reduced with lower physical activity. Proteins from each published study are indicated by different colors. Proteins overlapping between published datasets are shown in bold (5 elevated, 5 reduced), whereas proteins overlapping between the CP cohort and any of the published datasets are shown in bold and underlined (2).

**Supplementary Figure 5.**
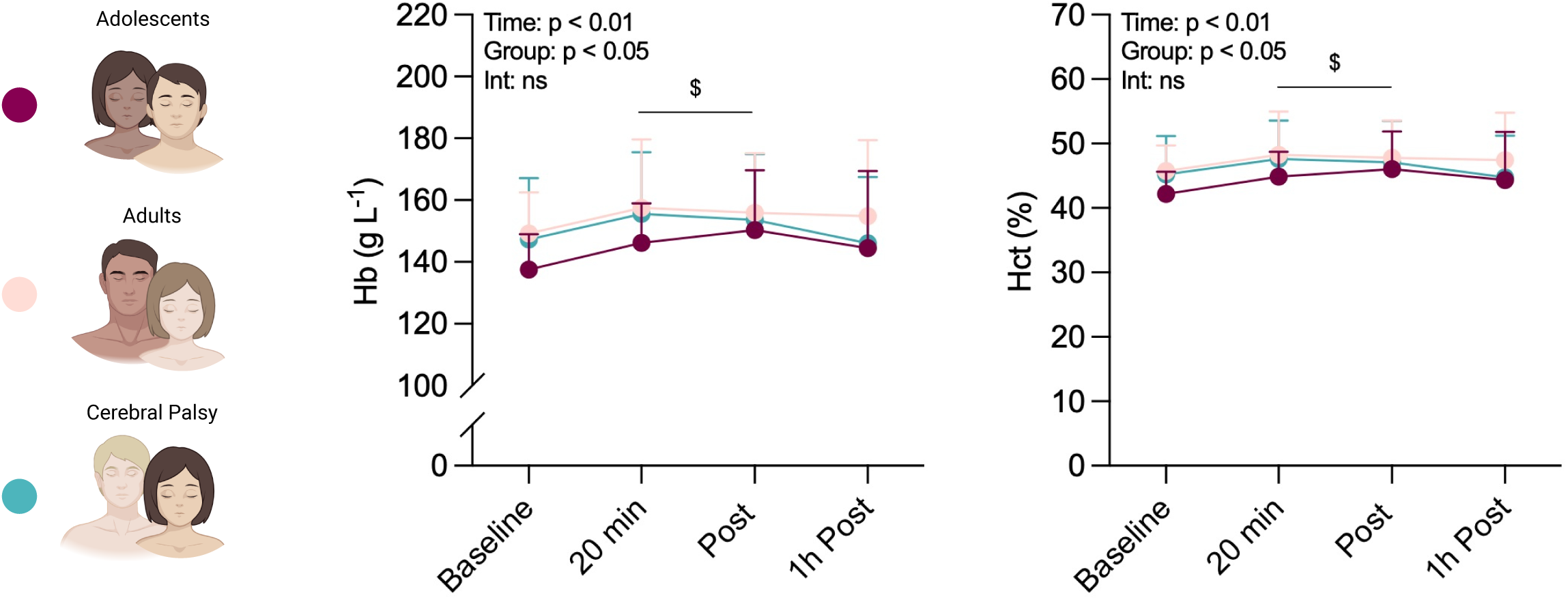
Additional blood gas changes in CP and TD individuals in response to an acute bout of endurance exercise. Hemoglobin (Hb) and hematocrit (Hct) at baseline, 20 minutes, immediately post-exercise, and one-hour post-exercise in TD adolescents, TD adults, and individuals with CP. Sample size: CP *n* = 12, TD adolescents *n* = 30, and TD adults *n* = 56. Changes in Hb and Hct over time and between groups were assessed using mixed-effects analysis; significant interactions were followed by post hoc comparisons using Dunnett’s test. $ indicates a significant difference compared to baseline (main effect of time). Statistical significance was defined as adjusted p < 0.05. Data are presented as mean ± SD for each group.

**Supplementary Figure 6A-B.**
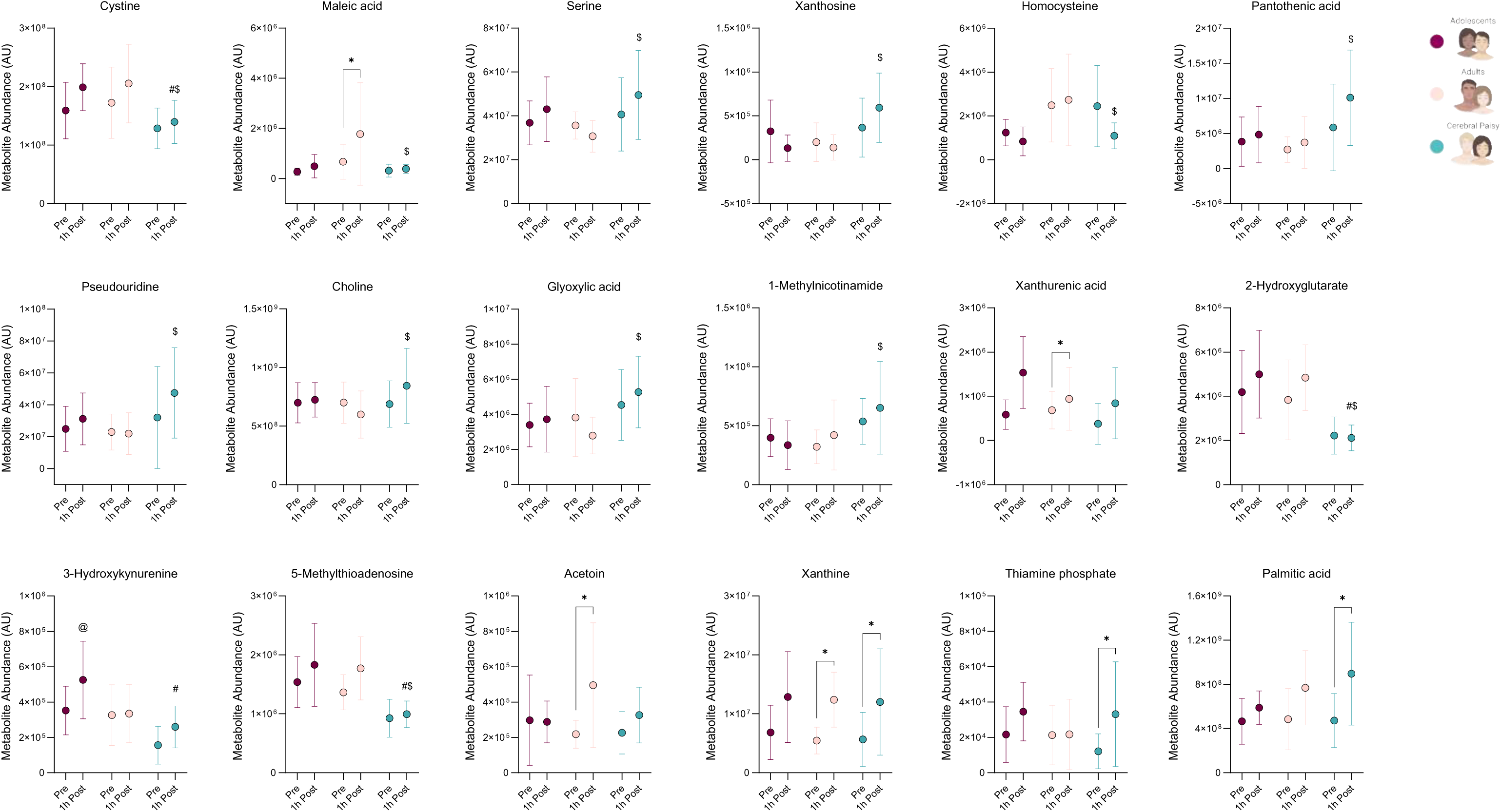
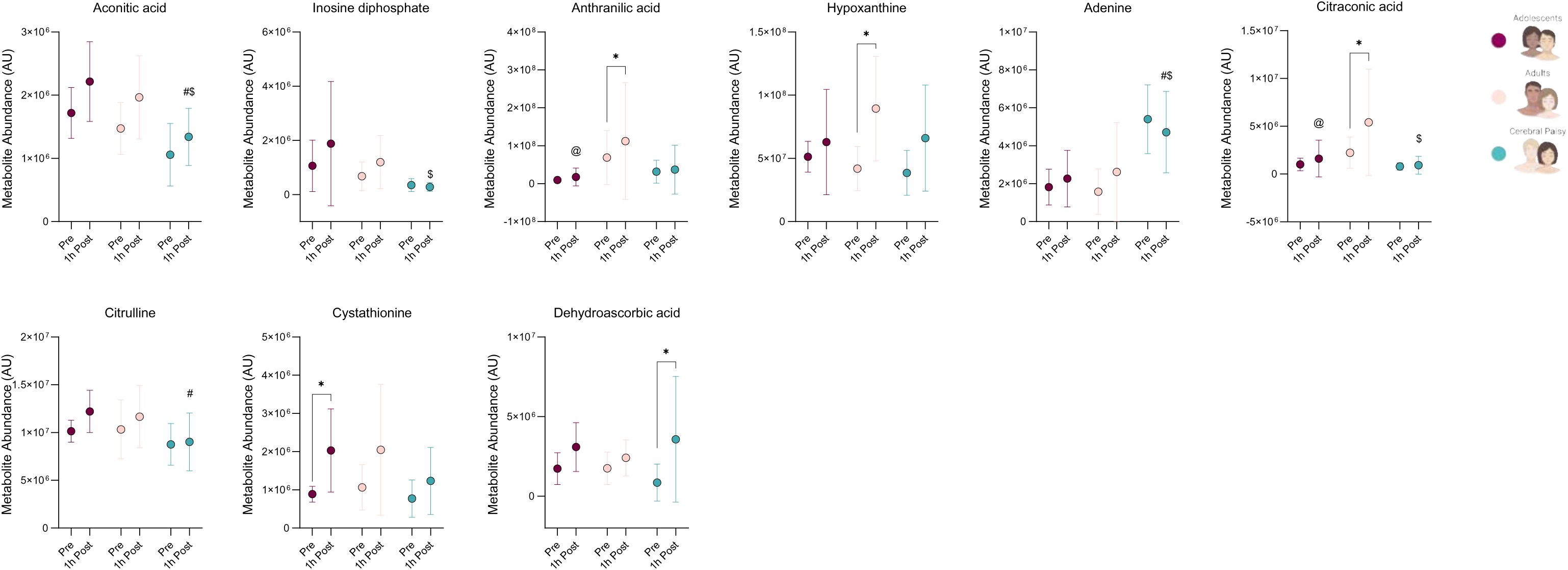
Additional plasma metabolite changes in CP and TD individuals in response to an acute bout of endurance exercise. Changes in metabolites before and after exercise in TD adolescents, TD adults, and CP adults. Sample size: CP *n* = 12, TD adolescents *n* = 13, and TD adults *n* = 17. Statistical significance of changes in metabolite levels in response to exercise was assessed using two-way ANOVA. P values were adjusted for multiple comparisons using Tukey’s post-hoc test. * indicates a significant pre- to post-exercise difference, $ indicates a significant difference between CP adults and TD adults at post-exercise, # indicates a significant difference between CP adults and TD adolescents at post-exercise, @ indicates a significant difference between TD adolescents and TD adults at post-exercise. Statistical significance was defined as adjusted p < 0.05. Data are presented as mean ± SD for each group.

**Supplementary Figure 7A-C.**
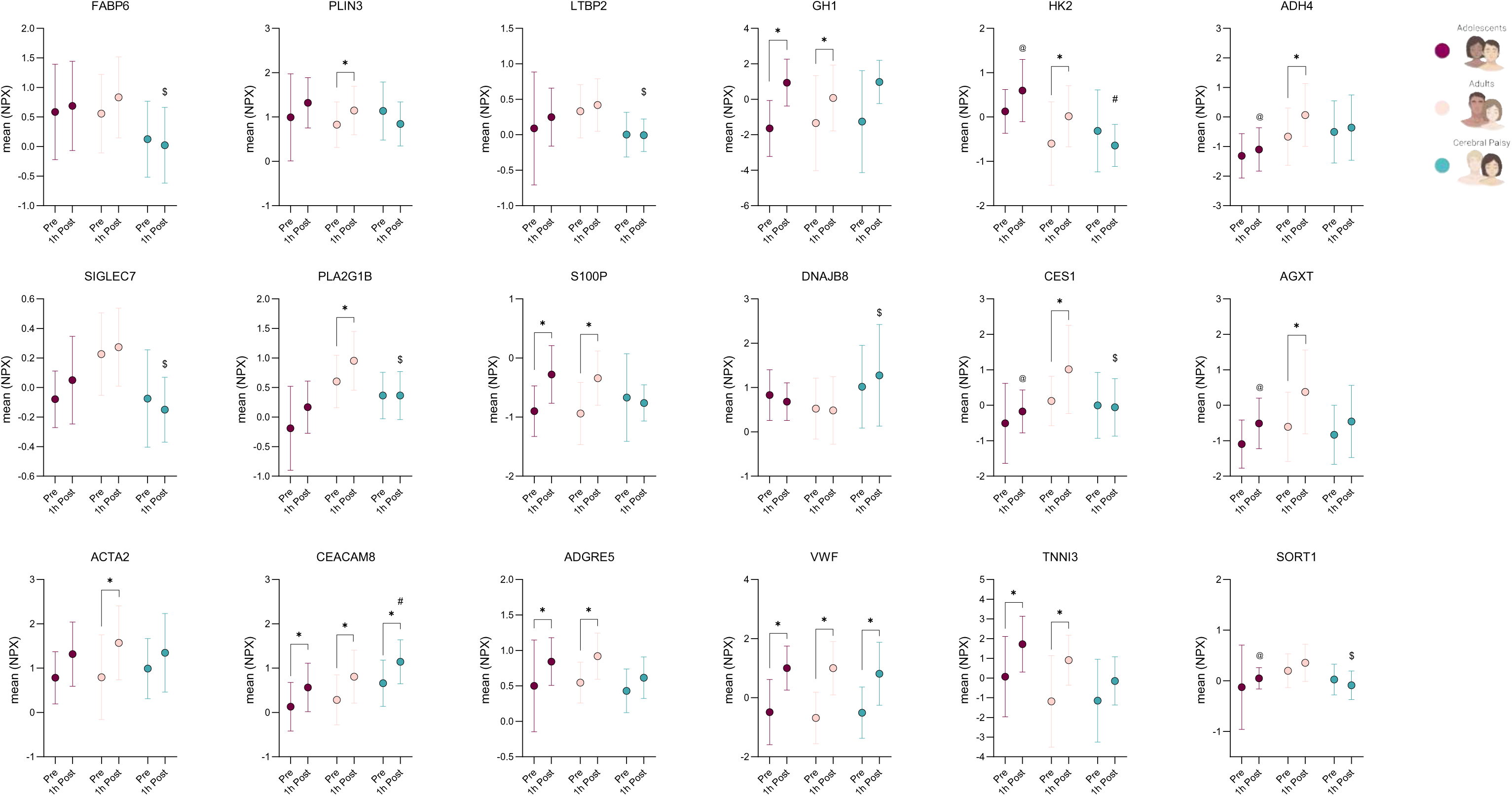
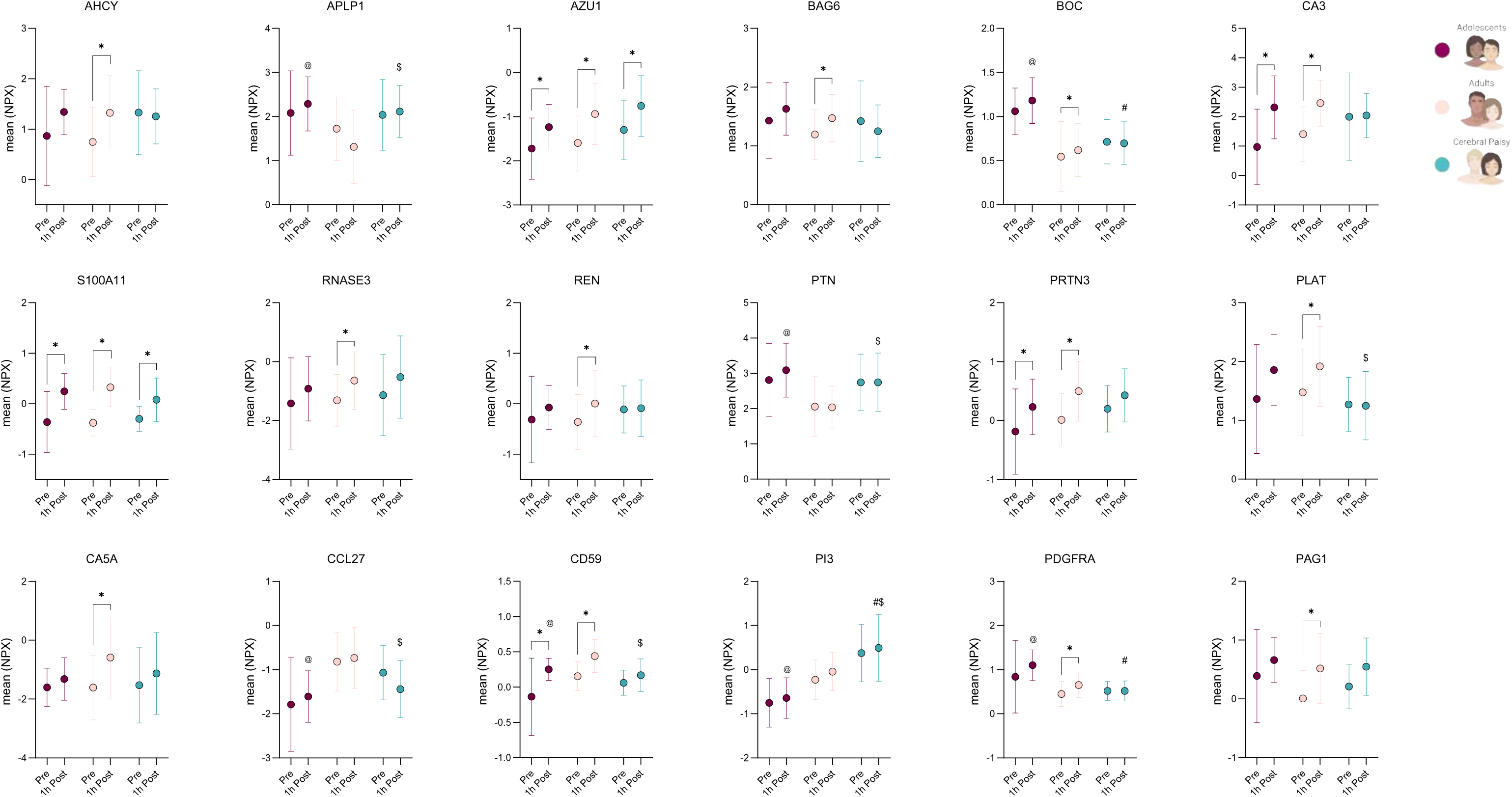
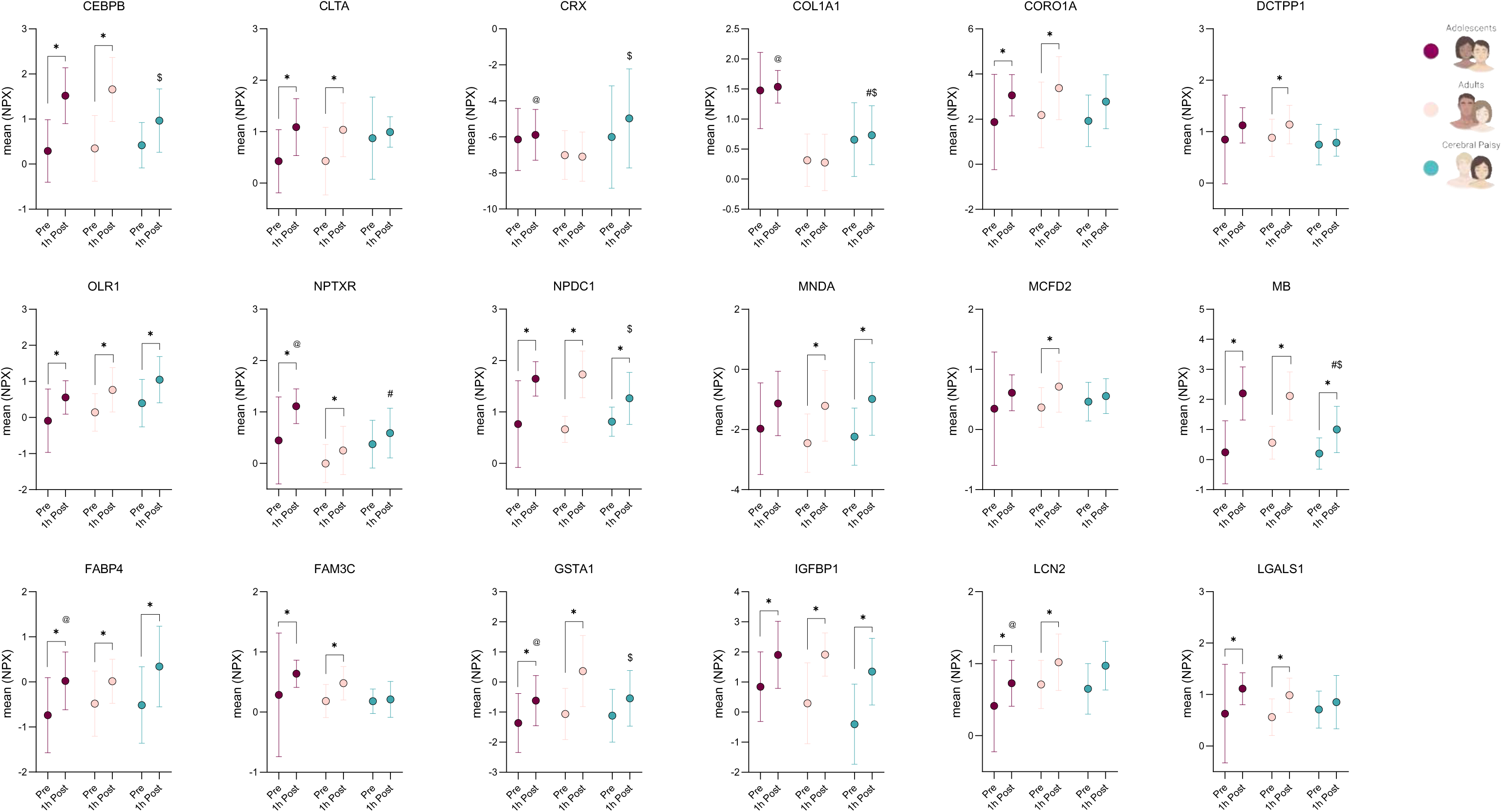
Additional plasma protein changes in CP and TD individuals in response to an acute bout of endurance exercise. Changes in proteins before and after exercise in TD adolescents, TD adults, and CP adults. Sample size: CP *n* = 15, TD adolescents *n* = 30, and TD adults *n* = 54. Statistical significance of changes in protein levels in response to exercise was assessed using two-way ANOVA. P values were adjusted for multiple comparisons using Tukey’s post-hoc test. * indicates a significant pre- to post-exercise difference, $ indicates a significant difference between CP adults and TD adults at post-exercise, # indicates a significant difference between CP adults and TD adolescents at post-exercise, @ indicates a significant difference between TD adolescents and TD adults at post-exercise. Statistical significance was defined as adjusted p < 0.05. Data are presented as mean ± SD for each group.

**Supplementary table 2.** Blood pressure and electrocardiographic parameters in CP and TD individuals. Data are presented as mean ± SD for normally distributed variables or as median (IQR) for non-normally distributed variables. Sample size: CP n = 12, TD adolescents n = 28, and TD adults n = 57. Between-group differences were assessed using one-way ANOVA with Tukey’s multiple comparisons test for normally distributed variables, or Kruskal-Wallis test with Dunn’s multiple comparisons test for non-normally distributed variables. # indicates a significant difference from TD adolescents, * indicates a significant difference from TD adults. Statistical significance was defined as adjusted p < 0.05. Abbreviations: IQR, interquartile range; SD; standard deviation.

|  | Typically developing<br><18 yrs | Typically developing<br>≥18 yrs | Cerebral palsy |
| --- | --- | --- | --- |
| Systolic blood pressure (mmHg) | 110.4 ± 9.5* | 119.1 ± 11.3<sup>#</sup> | 119.1 ± 15.9 |
| Diastolic blood pressure (mmHg) | 63.1 ± 8.4* | 75.3 ± 7.7<sup>#</sup> | 78.3 ± 12.6<sup>#</sup> |
| Heart rate (bpm) | 65.5 (57.0 – 71.0) | 62.0 (55.5 – 70.0) | 75.5 (64.5 – 101.8)* |
| PQ time (ms) | 136.1 ± 21.9* | 154.4 ± 23.3<sup>#</sup> | 141.1 ± 18.2 |
| QRS duration (ms) | 92.0 (88.0 – 96.8) | 96.0 (88.0 – 103.5) | 94.5 (86.0 – 102.8) |
| QTc (ms) | 416.8 ± 20.1 | 417.7 ± 18.3 | 419.1 ± 17.6 |

## Notes

### Competing Interest Statement

The authors have declared no competing interest.

